# Distinct Binding Mechanisms for Allosteric Sodium Ion In Cannabinoid Receptors

**DOI:** 10.1101/2021.04.07.438766

**Authors:** Soumajit Dutta, Balaji Selvam, Diwakar Shukla

## Abstract

The therapeutical potential of Cannabinoid receptors is not fully explored due to psychoactive side-effects and lack of selectivity associated with the orthosteric ligands. Allosteric modulators have the potential to become selective therapeutics for cannabinoid receptors. Biochemical experiments have shown the effects of the allosteric Na^+^ binding on cannabinoid receptor activity. However, the Na^+^ coordination site, and binding pathway are still unknown. Here, we perform molecular dynamic simulations to explore Na^+^ binding in the cannabinoid receptors, CB_1_ and CB_2_. Simulations reveal that Na^+^ binds to the primary binding site from different extracellular sites for CB_1_ and CB_2_. A distinct secondary Na^+^ coordinate site is identified that is not present in CB_2_. Furthermore, simulations also show that intracellular Na^+^ could bind to the Na^+^ binding site in CB_1_. Constructed Markov state models show that the standard free energy of Na^+^ binding is similar to the previously calculated free energy for other class A GPCRs.

## Introduction

Cannabinoid receptors are part of the endocannabinoid system, which control cellular homeostasis by intracellular signal transduction (*1*). In the last decade of the twentieth century, cannabinoid receptor 1 (CB_1_) and 2 (CB_2_) were discovered as a target of cannabinoid compounds, the major constituents of marijuana (*2, 3*). CB_1_ and CB_2_ belong to the class A GPCRs (*4*), the largest subfamily of GPCR proteins. CB_1_ receptor, which is majorly expressed in the central and peripheral nervous system, has therapeutic potential for pain, obesity, and addiction (*1, 5*–*9*). On the other hand, CB_2_ receptors are expressed in human immune cells and peripheral tissues, and can be targeted for inflammatory, fibrotic diseases (*10, 11*). However, orthosteric ligands of these receptors are not available on the market as therapeutics due to lack of selectivity and over-stimulation effects. The lack of selectivity can be explained by the similarity of structure and sequence between the two cananbinoid receptors (*12, 13*). Furthermore, orthosteric agonists (e.g., fubinaca) and antagonists (e.g., rimonabant) of CB_1_ lead to overstimulated response (*9, 14*) and harmful side-effects like anxiety, depression (*15*). Therefore, to develop a therapeutic drug, an allosteric modulator can be a potential option due to two key reasons (*16, 17*). First, an allosteric site of a receptor is generally less conserved as compared to the orthosteric binding site. Hence, allosteric ligands can be more selective than orthosteric ligands (*16*). For instance, the allosteric ligand ORG27569 regulates the binding affinity of an orthosteric agonist CP55940 for CB_1_, but not for CB_2_ (*18*). Second, allosteric ligands can only regulate receptor function in the presence of orthosteric ligands and have ceiling efficacy controlling orthosteric ligand function (*16, 17*). Therefore, allosteric ligands could avoid over-stimulation effects. One such allosteric site for CB_1_ and CB_2_ is the Na^+^ ion binding site coordinated by conserved D^2.50^ and surrounding hydrophilic and hydrophobic residues (*19*–*21*). Na^+^ acts as a conserved negative allosteric modulator (NAM) for class A GPCRs. This Na^+^ binding site has been targeted for potential NAM drug design, which can mimic the effect of Na^+^. For instance, amiloride and its derivative have been shown to compete with Na^+^ for binding in the allosteric site for different class A GPCRs (*22, 23*). Therefore, elucidation of Na^+^ binding site and binding pathways for CB_1_ and CB_2_ are essential for designing a therapeutically selective allosteric modulator drug which can target this site.

Although biochemical experimental evidence showed that Na^+^ acts as a NAM for CB_1_ and CB_2_, none of the x-ray crystal or cryo-EM structures captured Na^+^ in its putative binding site (*5*–*9, 11, 24, 25*). Therefore, the Na^+^ co-ordination site is unknown for both these receptors. Previous structural studies have also shown that Na^+^ co-ordination site shifts towards the intracellular site by one helical turn due to mutation of conserved N^7.49^ to D^7.49^ (*23, 26*). Superposition of the inactive structure of CB_1_ (PDB ID: 5TGZ (*5*)) and CB_2_ (PDB ID: 5ZTY (*11*)) reveals two key structural and sequence changes (Figure S1). First, The conserved S^3.35^ residue in CB_1_ is mutated into T^3.35^ in CB_2_. Due to this mutation, T^3.35^ moves away from conserved the D^2.50^ in CB_2_. Second, Y^3.35^ sidechain in the NPxxY motif in CB_1_ points upward towards to Na^+^ binding site while in CB_2_ it points away from the Na^+^ binding site towards TM5 (Figure S1). Therefore, we hypothesize that these changes in sequence and structure between receptor’s allosteric Na^+^ binding site lead to distinct Na^+^ coordination sites which can significantly affect their drug pharamacophore preferences.

It is hard to determine the Na^+^ binding pathway experimentally because the other transporters and Na^+^ channels present in the cell can also affect the measurement of Na^+^ flux or voltage difference between the two sides of the membrane. Hence, computational studies using molecular dynamics (MD) has been crucial to discover binding pathways for Na^+^ (*27*–*33*) and understand conformational equilibrium of GPCRs (*34, 35*). In particular, Selvam et al. has simulated the Na^+^ binding pathways for 24 families of GPCRs to conclude that class A GPCRs follow a universal ion-binding mechanism (*33*). This study also showed that Na^+^ could enter the binding site from both extracellular and intracellular region for different GPCRs. Comparative study on Na^+^ binding for different opioid receptors has shown that Na^+^ enters from different extracellular regions for *µ, κ, δ*-OR due to variation of the negatively charged residues in the loop regions of the receptors (*27*). Structural comparison of both cannabinoid receptors shows that the N-loop moves towards the orthosteric binding pocket and has acidic residues for CB_1_ while for CB_2_ it floats outside the orthosteric pocket (Figures 1A and 1B). N-loop positioning can significantly affect the Na^+^ binding pathways for the receptors. Therefore, the Na^+^ binding pathway for CB_1_ and CB_2_ may also be different.

**Figure 1:**
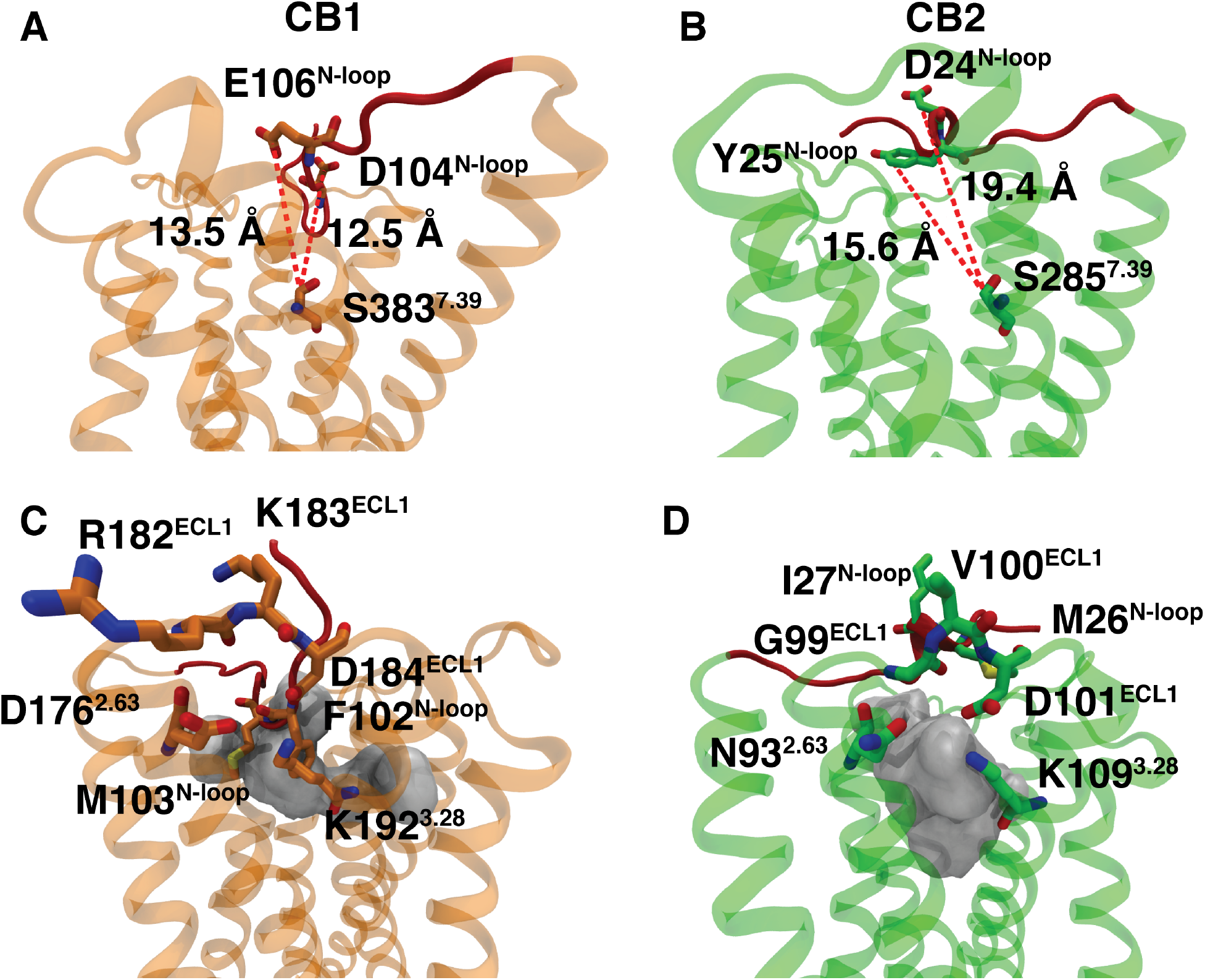
Extracellular Na^+^ binding site comparison. The distances between polar residue in orthosteric binding pocket and polar residues in N-loop for inactive CB_1_ (PDB ID: 5TGZ (*5*)) (A) and CB_2_ (PDB ID: 5ZTY (*11*)) (B). The accessibility of orthosteric binding pocket volume from ECL1 site for inactive CB_1_ (PDB ID: 5TGZ (*5*)) (C) and CB_2_ (PDB ID: 5ZTY (*11*)) (D). Proteins are shown with cartoon representation (CB_1_: orange, CB_2_: green). N-loops are colored as red. Important residues are represented as sticks. Measured distances are shown as dotted line. Pocket volume was measured using Fpocket (*36*).

To resolve Na^+^ binding for cannabinoid receptors, we run atomic scale extensive molecular dynamics simulations to predict and compare the Na^+^ binding site and pathway for each receptor. To determine the thermodynamics and kinetics of the binding process, Markov state models (MSM) are built using simulation data. Our results show that Na^+^ binds via a distinct pathway from extracellular site to each of the receptors. For CB_1_, Na^+^ binds with the help of negatively charged residues of downward N-loop inside orthosteric binding pocket; alternatively, Na^+^ binds for CB_2_ from the gap formed between TM1, TM2 and ECL1. Furthermore, we find an additional secondary binding site for Na^+^ between TM1, TM2, and TM7 for CB_1_. This site does not exist in CB_2_. Our results also reveal that Na^+^ can bind to the secondary binding site from the intracellular region. By determining new Na^+^ binding site and identifying differences in the binding pathway between CB_1_ and CB_2_, this study will aid in design of new allosteric drugs for cannabinoid receptors.

## Results

### Extracellular binding of Na^+^ to cannabinoid receptors

Comparing Na^+^ binding simulation data of CB_1_ and CB_2_ reveals that Na^+^ can bind to receptors from extracellular region, as shown in previous simulation studies for other GPCR proteins (*27, 33*). However, the exact extracellular Na^+^ binding site varies for CB_1_ and CB_2_ due to the structural and sequence variation between the receptors. For CB_1_, Na^+^ binds with the assistance of negatively charged residues in N-loop (D104^N*−*loop^). In the inactive structure of CB_1_, the N-loop moves inside the orthosteric pocket which brings the D104^N*−*loop^ adjacent to conserved polar residue S383^7.39^ and backbone carbonyl group of A257^7.36^ (Figure 1A). D104^N*−*loop^ assists the movement of the Na^+^ from the extracellular solvent layer to the orthosteric binding pocket (Figure S2A). During activation of CB_1_, this N-loop moves away from the pocket which may hinder the Na^+^ binding from the extracellular side (*6*). Therefore, downward movement of N-loop may play a major role in stabilizing inactive states of CB_1_ by aiding the binding of negative allosteric modulator Na^+^. Although, CB_2_ also has negatively charged (D24^N*−*loop^) and polar residue (Y25^N*−*loop^) in the N-loop, major conformational change in the N-loop region is not observed between active and inactive structure of CB_2_ as compared to CB_1_(*24, 25*). Therefore, D24^N*−*loop^ and Y25^N*−*loop^ remain outside the pocket and interact with Na^+^ ion from extracellular region for 14 *±* 0.2% of the simulation time but cannot transfer the ion to the orthosteric pocket (Figures 1B and S3A).

We observe that the Na^+^ binds from the gap between TM2, TM3, and ECL1 for CB_2_. For CB_2_, this region is connected to the orthosteric pocket as shown in Figure 1D. Negatively charged D101^ECL1^, polar N93^2.63^ and backbone carbonyl group of G99^ECL1^ helps Na^+^ ion to bind and move to the orthosteric pocket of the CB_2_ (Figure S2B). Previous studies have shown that D101^ECL1^ residue is also important for binding of other ligands to CB_2_ (*37*). CB_1_ also have conserved D184^ECL1^ residue in this position but the residue is surrounded by bulkier and positively charged R182^ECL1^ and K183^ECL1^ (Figure 1C). These positively charged residues block the access of D184^ECL1^ to the extracellular Na^+^. In the simulated ensemble, only 1.6 *±* 0.3% MSM weighted frames of CB_1_ have Na^+^ bound to D184^ECL1^, whereas, in CB_2_, 8.0 *±* 0.5% frames have Na^+^ bound to D101^ECL1^ (Figure S3B). Furthermore, the path from the ECL1 to othrosteric pocket is hindered by downward hydrophobic N-loop residues for CB_1_ as shown in figure (Figure 1C). Therefore, the structural and sequence variation in the N-loop and ECL1 site leads to different binding sites for Na^+^ ion in the cannabinoid receptors.

After recognition by the extracellular binding site, Na^+^ moves to the orthosteric binding pocket for CB_1_ and CB_2_. The orthosteric pocket of these receptors mostly consist of bulky hydrophobic residues except for S383^7.39^ (CB_1_) or S285^7.39^ (CB_2_). MSM-weighted free energy landscape projection of y and z coordinate of Na^+^ with respect to D163^2.50^ of CB_1_ shows that the activation energy required to cross the free energy barrier is *∼* 3 *±* 0.2 kcal/mol (Figures S4A and S5A). Along the pathway, the Na^+^ ion interacts with residues from transmembrane TM2, TM3, and TM7 (Figure S4C). To evaluate important residue movements inside the orthosteric pocket, we perform time independent component analysis (tiCA) on our simulation data. It reveals that the movement of F170^2.57^ is crucial and it is one of the slowest process during the Na^+^ binding (Figure S6C). MSM weighted free energy landscape shows that ensemble average distance between F170^2.57^ (C*γ*) and V196^3.32^ (C*γ*) is comparatively higher when Na^+^ is in orthosteric binding pocket (9.5 °A) compared to when Na^+^ is in bulk (7.1 °A) (Figures 2A, S7A and 2C). Due to this movement of conserved residues and the flexible N-loop, pore tunnel of radius 2.1 *±* 0.1 Å is created inside the binding pocket (Figure S8) for CB_1_. For CB_2_, similar interactions are observed in orthosteric binding pocket. The activation barrier to move the orthosteric pocket from ECL1 binding site is close to *∼* 3.5*±*0.2 kcal/mol (Figures S4B, and S5B). Conserved F87^2.57^ residue in the similar position as CB_1_ is also found to be crucial for Na^+^ movement in CB_2_ (Figures S6B). In this case, the average distance between F87^2.57^ and V113^3.32^ increases by 2.4 Å when Na^+^ is bound inside the pocket (Figures 2B, S7B and 2D). The radius of the pore tunnel generated by this movement is 2.6 Å (Figure S8). Therefore, these residue movements facilitate the Na^+^ to cross the hydrophobic barrier in the orthosteric pocket and bind to Na^+^ binding pocket.

**Figure 2:**
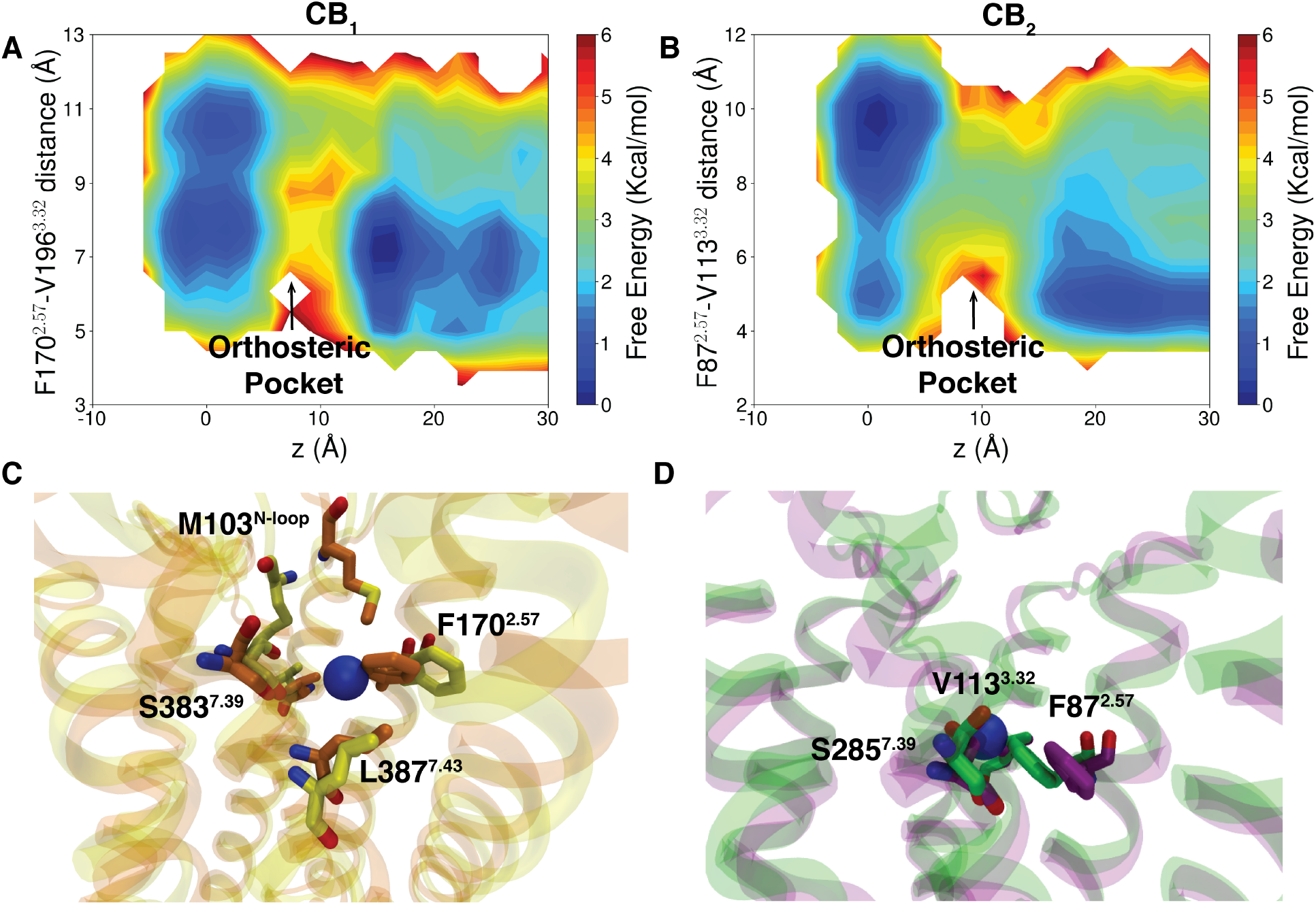
Important residue movement within the orthosteric binding site facilitating Na^+^binding. (A) shows MSM weighted free energy landscape projected along z component distance of Na^+^ from D163^2.50^ (O*δ*1) and F170^2.57^ (C*γ*) - V196^3.32^ (C*γ*) distance for CB_1_. (B) shows MSM weighted free energy landscape projected along z component distance of Na^+^ from D80^2.50^ (O*δ*1) and F87^2.57^ (C*γ*) - V113^3.32^ (C*γ*) distance for CB_2_. (C) and (D) represent superposition of inactive structure and MD snapshot where Na^+^bound in the orthosteric pocket for CB_1_ and CB_2_. Proteins are represented as cartoon (Inactive CB_1_: orange, inactive CB_2_: green, CB_1_ MD snapshot: yellow, CB_2_ MD snapshot: violet). Important residues are represented as sticks. Na^+^s are represented as VDW representation (color: blue).

### Na^+^ binding site for CB_1_ and CB_2_

From the orthosteric binding site, Na^+^ moves to the primary Na^+^ binding site. To calculate the timescale for the transition between orthosteric pocket and Na^+^ binding site, we measure the mean free passage time using transition path theory (TPT). The timescale for transition is similar for both the receptor (Figures 3A, 3B, and S9). For CB_1_, this transition happens in 3.0 *±* 1.6*µs* whereas for CB_2_ it takes 0.5 *±* 0.4*µs*. In the primary binding site, Na^+^ is co-ordinated by D^2.50^, S^3.39^, and N^7.45^, which is similar to other class A GPCRs from the same branch (*α* branch) (Figures S10A, S10B, S11A, and S11B). However, superimposing the MD snapshots of CB_1_ and CB_2_ with other branches of class A (*δ* (PAR1) and *γ*(*δ − OR*)) shows that Na^+^ coordination site differs slightly. For PAR1, Na^+^ binds towards intracellular side as compared to CB_1_ and CB_2_ and interacts with D^7.49^ instead on N^7.45^ (Figures S10C and S11C). In case of *δ − OR*, Na^+^ binds closer to TM3 as compared to CB_1_ and CB_2_ (Figures S10D and S11D).

**Figure 3:**
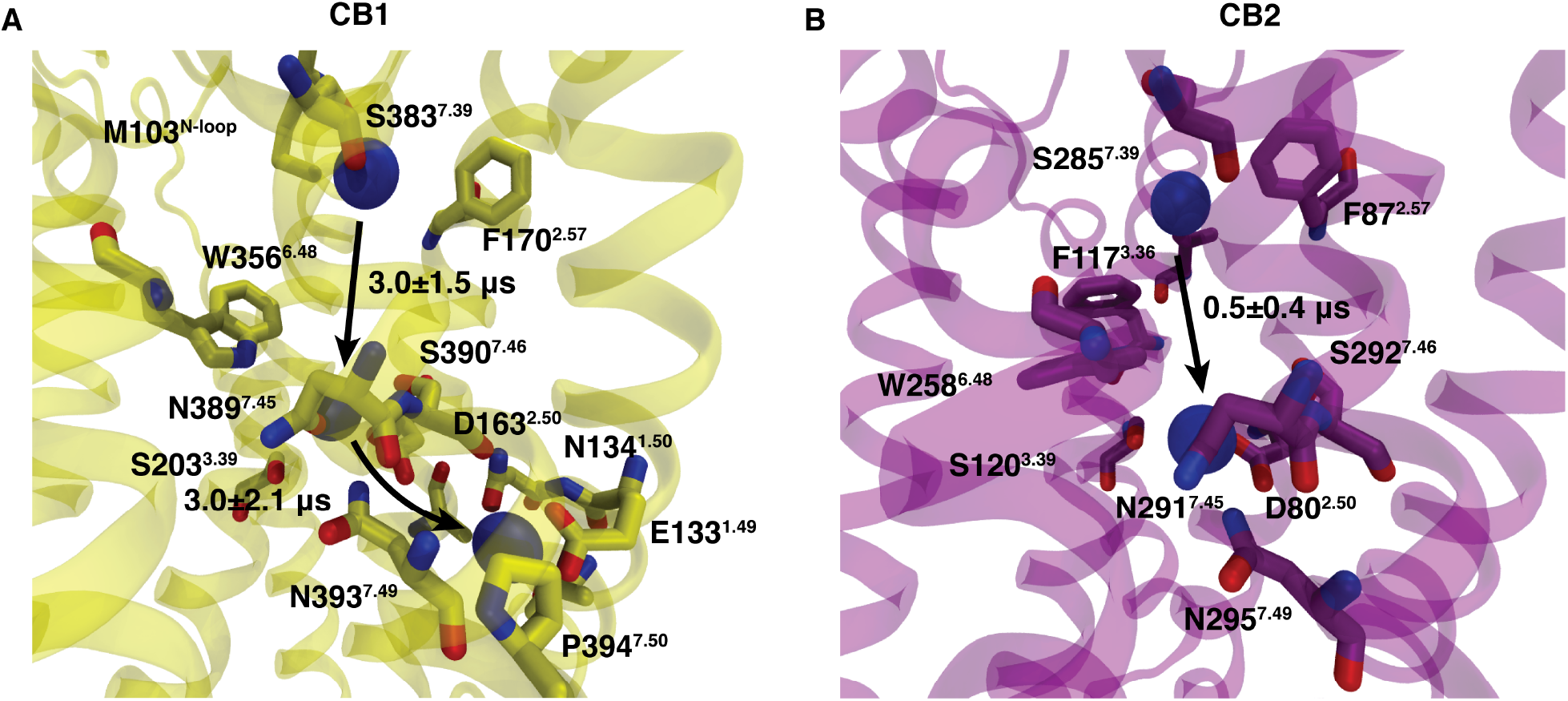
Binding of Na^+^ from orthosteric binding site to the Na^+^ binding pocket. (A) Representation of Na^+^ in different positions in CB_1_ (orthosteric binding site, primary Na^+^binding site, secondary Na^+^ binding site). (B) Representation of Na^+^ in different positions in CB_2_ (orthosteric binding site, primary Na^+^ binding site). Arrows represent the direction of Na^+^ movement. Important residues are represented as sticks. Proteins are shown as Cartoon (CB_1_: yellow, CB_2_: magenta).

On top of the primary binding site, we discover a secondary binding site for CB_1_ between TM1, TM2 and TM7 which does not exist for CB_2_ (Figures 3A, S4A, and S4B). To characterize this newly found site, MSM weighted free energy landscape between Y153^2.40^-Y397^7.53^ and D163^2.50^-N393^7.49^ residue distance were estimated (Figure 4A). The two stable states could be seen in the landscape. In the state I, Y397^7.53^ faces towards the binding site and N393^7.49^ remains close to D163^2.50^ (Figure 4C). In state II, Y397^7.53^ moves away from the Na^+^ binding pocket. Due to the this movement, N393^7.49^ moves away from D163^2.50^. This larger distance between TM2 and TM7 allows Na^+^ to migrate to the secondary binding site. This binding site is surrounded by residues polar (N134^1.50^, N393^7.49^) and negatively charged residues (D163^2.50^, E133^1.49^) (Figure 3A). Thermodynamic calculations show that the activation barrier for this transition is *∼* 1.5 *±* 0.3 kcal/mol (Figure S4A and S12B). Moreover, kinetic calculations with TPT analysis show that forward and backward transition between the primary and secondary binding site happens within the microsecond timescale (Figure S13). Therefore, thermodynamically and kinetically Na^+^ binding is feasible for either binding site in CB_1_.

**Figure 4:**
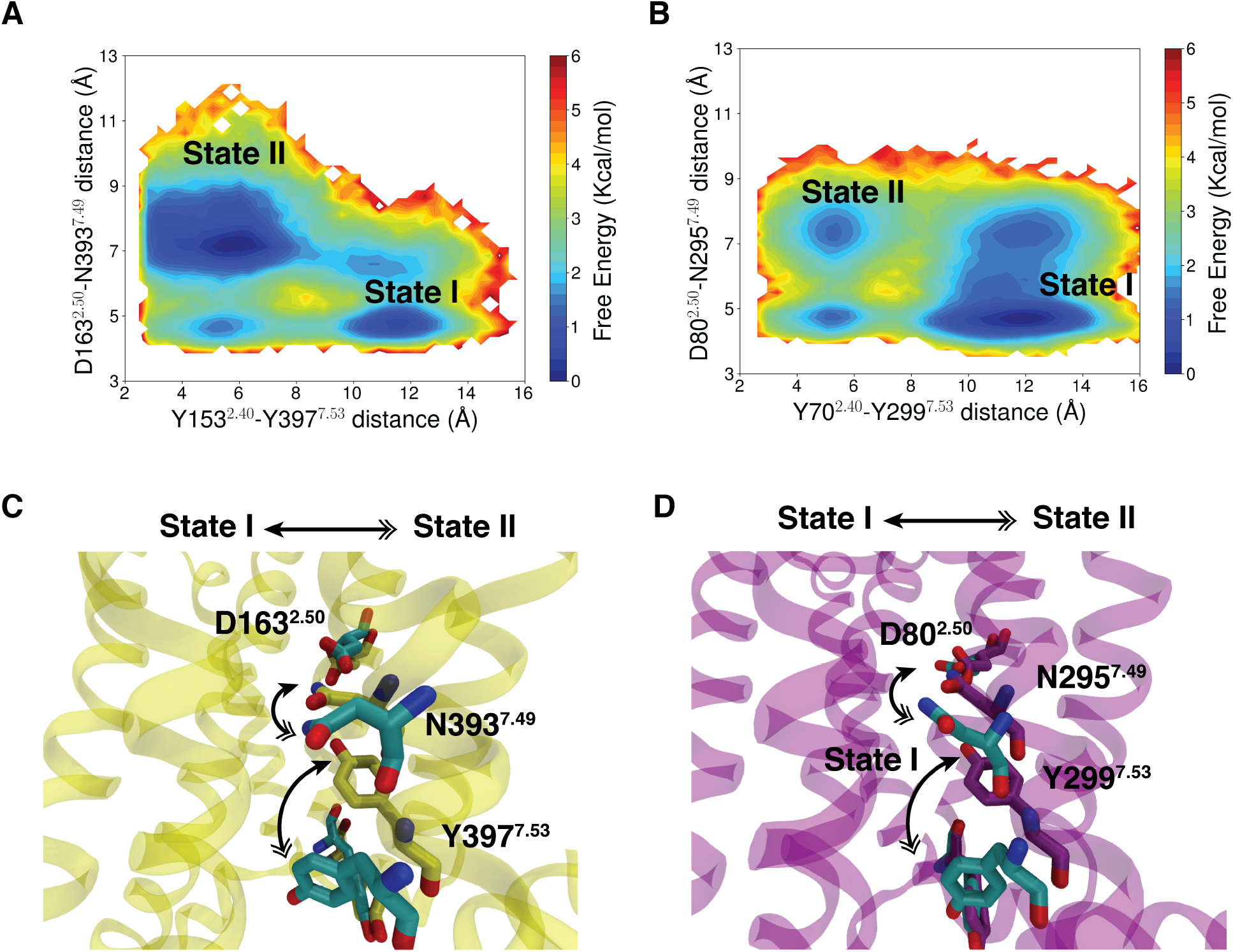
TM7 movement leads to larger distance between polar residues in the Na^+^ binding pocket. (A) MSM weighted free energy landscape projected along Y153^2.40^ (O*η*) - Y397^7.53^ (O*η*) distance and D163^2.50^ (C*γ*) - N393^7.49^ (C*γ*) distance for CB_1_. (B) MSM weighted free energy landscape projected along Y70^2.40^ (O*η*)-Y299^7.53^ (O*η*) distance and D80^2.50^ (C*γ*) - N295^7.49^ (C*γ*) distance for CB_2_. (C) shows the residue movement between state I and state II.

Similarly for CB_2_, we observe both the states in MSM weighted free energy landscape estimated using distance betweeen Y70^2.40^-Y299^7.53^ and D80^2.50^-N295^7.49^ residue pairs (Figures 4B and S12B). However, state II has less population in CB_2_ compared to CB_1_ (Figure S14). Therefore, TM2 remains close TM7 and Na^+^ has less probability to move to the secondary binding site. Furthermore, comparison between volume distribution of secondary binding site for CB_1_ and CB_2_ shows that CB_1_ has larger volume as compared to CB_2_ (Figure S15). To understand the structural reasons behind this volume change, the distance distribution between conserved residues in TM1 (N^1.50^)and TM7(N^7.49^) was estimated. It shows that TM1 and TM7 are more distant from one another in CB_1_ as compared to CB_2_ (Figure S16A). Lower flexibility in TM1 for CB_2_ may emerge due to the steric hindrance from bulky F^2.51^ residue in TM2, which blocks the outward movement of TM1. Superimposition of the both inactive structure show that CB_1_ has smaller V^2.51^ residue in the same position (Figure S16B). Therefore, lower flexibility of TM1 and TM7 explains the absence of secondary binding site in CB_2_.

### Intracellular Na^+^binding for CB_1_

Selvam et al. has shown that Na^+^ could bind from the intracellular side in GPCRs (*33*). For CB_1_, the similar phenomena is observed. Na^+^ binds from the intracellular side between the gap of TM1 and TM7 (Figures 5A, 5C, and S17A). Negatively charged E133^1.49^ and polar T391^7.47^ helps Na^+^ moves inside the secondary binding site. MSM-weighted free energy landscape shows that side-chains of these two residues come close to drive the Na^+^ inside the secondary binding pocket (Figures 5A and 5C). Previous studies have also shown that E133^1.49^ acts as a potentially allosteric binding side for CB_1_ (*38, 39*).

**Figure 5:**
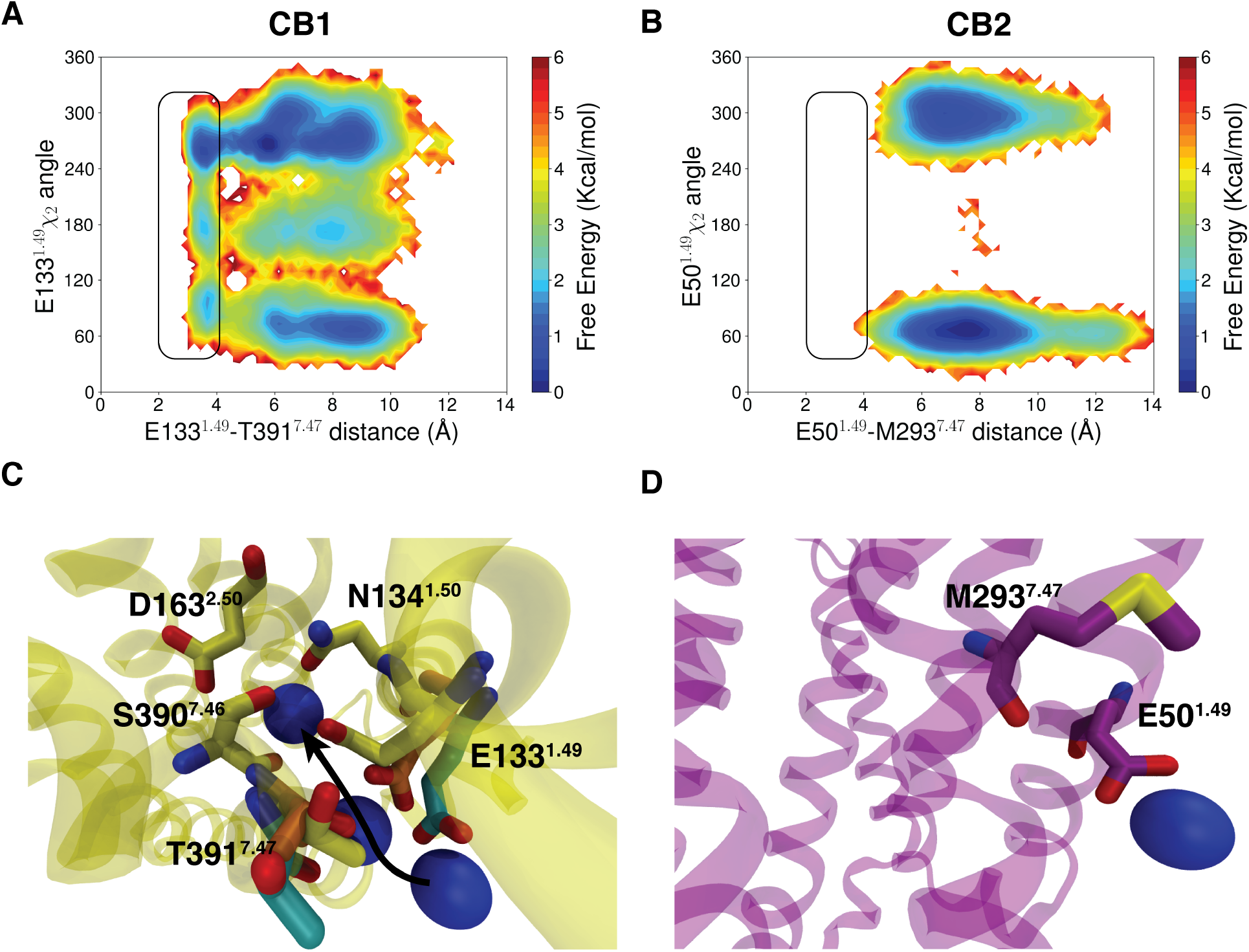
Na^+^binding from intracellular region. (A) MSM weighted free energy landscape between E133^1.49^ (C*δ*) -T391^7.47^ (O*γ*) distance and E133^1.49^ sidechain *χ*_2_ angle for CB_1_. (B) MSM weighted free energy landscape between E50^1.49^ (C*δ*) -M293^7.47^ (S*δ*) distance and E50^1.49^ sidechain *χ*_2_ angle for CB_2_. (C) shows intracellular binding of Na^+^ for CB_1_. Arrow is representing the pathway for Na^+^binding. E133^1.49^, T391^7.47^ are colored in differently to show different stages of binding (Na^+^ in intracellular region: cyan, Transition state: orange, Na^+^ bound to secondary binding site: yellow). Proteins are represented as Cartoon (CB_1_: yellow, CB_2_: voilet). Important residues are represented as sticks.

Although CB_2_ has conserved E50^1.49^ and Na^+^ occupancy of the E50^1.49^ site is 14.6*±*1.2%, intracellular binding for CB_2_ is not detected in our simulations (Figure S3C). There can be two explanations for this phenomenon. CB_2_ has hydrophobic M293^7.47^ instead of polar T391^7.47^ residue that can hinder the Na^+^ binding from inside the receptor. Furthermore, backbones of TM1 and TM7 remain close together as discussed in the previous section (Figure S18A). Therefore, E50^1.49^ side chain cannot enter the secondary binding site to facilitate Na^+^ binding as shown in Figures 5B and 5D. Therefore, these structural and sequence changes in TM1 and TM7 region explain absence of intracellular Na^+^ binding for CB_2_.

We also compare the CB_1_ intracellular binding site with the GPCRs where intracellular Na^+^ binding was established before such as PAR1 and P2Y12. It shows that PAR1 has hydrophobic residue in place of E^1.49^. However it has P^1.48^ close to intracellular binding site which gives TM1 more flexibility to move (Figure S18B). In case of P2Y12, we observe a polar residue T^1.49^ in the same position (Figure S18C). Therefore, flexibility in TM1 and TM7 and polar residues in the 1.49 position may be deterministic factor for intracellular Na^+^ binding.

## Conclusions

In this study we compare the Na^+^ binding processes for two cannabinoid receptors. Although these two receptors share 44% sequence similarity and orthosterically bind similar classes of ligands, we observe clear differences in their respective Na^+^ binding sites and pathways. Kinetically extracellular Na^+^ binding is faster for CB_2_ compared to CB_1_ (Figures 6A, 6B). Whereas, thermodynamics calculations show opposite trend for extracellular binding of Na^+^ (Figure S19A). Standard binding free energy for extracellular Na^+^ binding for CB_1_ and CB_2_ is in the same range as previously calculated free energies (2-5 kcal/mol) for other class A GPCR (*33*). We also observe intracellular binding of Na^+^ in CB_1_ and the standard binding free energy for extracellular and intracellular binding are energetically similar. However, kinetically intracellular binding of CB_1_ is more accessible (Figures 6A, and S19B). Comparison with previously reported intracellular binding of Na^+^ shows that TM1 and TM7 flexibility and polar residue at 1.49 position is required for Na^+^ to bind from intracellular direction. Future studies may reveal intracellular Na^+^ binding for other class A GPCRs.

**Figure 6:**
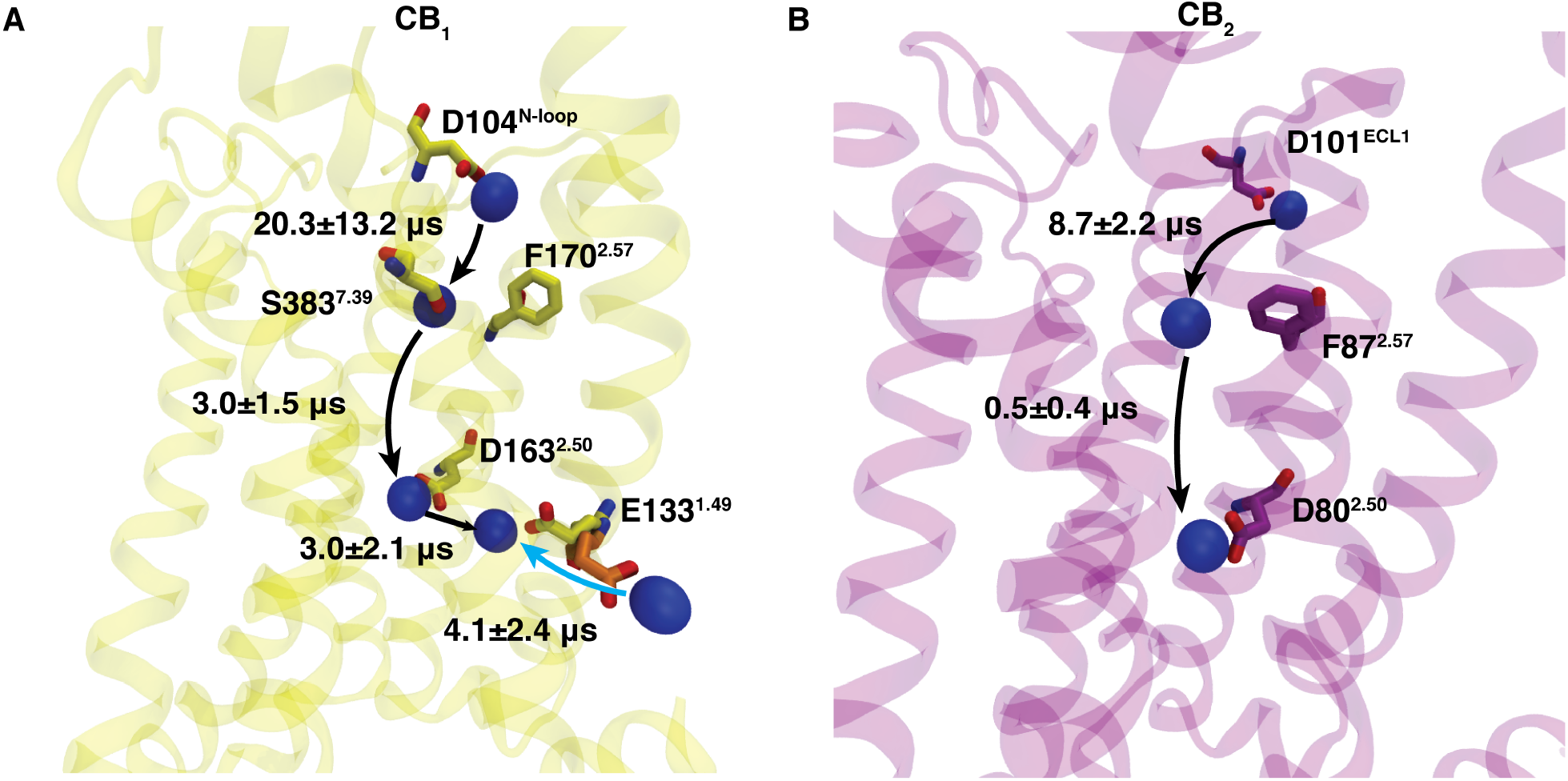
Overall pathway for Na^+^ binding for CB_1_ (A) and CB_2_ (B). Timescales for jump between two important steps are shown by TPT analysis. E133^1.49^ residues are shown in two different color to represent two different states (Na^+^ bound to secondary binding site: yellow, Na^+^ in the intracellular region: orange).

We also observe a secondary Na^+^ binding site for CB_1_ which has not been previously reported. This site is accessible from both intracellular and extracellular side. The inward movement of E133^1.49^ stabilize Na^+^ in that position (Figures 6A) which has been previously reported as allosteric binding site(*38, 39*). Although Na^+^ binding is conserved for all class A GPCRs, large differences may exist in Na^+^ binding co-ordination sites and binding pathway due to evolutionary divergence in GPCR sequence. These changes in Na^+^ binding could be exploited for the design of allosteric modulators of GPCRs.

## Methods

### System Preperation

The crystal structures of inactive CB_1_(PDB ID: 5TGZ(*5*)) and CB_2_ (PDB ID: 5ZTY (*11*)) were used as a starting structure for the simulation. Ligands, other non-protein molecules and the stabilizing fusion parter between TM5 and TM6 were removed from the crystal structures. Thermostabilized residues in protein crystal structure were mutated back according to the original protein sequence (*5, 11*). Hydrogen atoms were added to the system using the reduce command in AMBER tools. Truncated N-loop, C-loop and unconnected TM5 and TM6 were neutralized by adding acetyl (ACE) and methylamide (NME) capping groups. To embed proteins within a membrane environment and to solvate the extracellular and intracellular region, CHARMM-GUI software was used (*40*). We employ POPC membrane layer for our simulation. A physiological salt (Na^+^ and Cl-) concentration of 150 mM and TIP3P water model were used to solvate the system. Lastly, system parameterization was done using AMBER ff14SB and Lipid17 forcefield ((*41, 42*)).

### Simulation Details

AMBER18 package was used to run MD simulations on BlueWaters Supercomputer at National Center for Supercomputing Applications(*43*). Before proceeding to the production stage of simulation, system was minimized and equilibrated. System minimization was done using gradient descent and conjugate gradient algorithm for 15000 steps. Minimized system was heated in NVT ensemble to increase the temperature of the system gradually from 0 to 10K and 10K to 300K. Each heating step is performed for 1 ns. To maintain the pressure of heated system at 1 bar, NPT ensemble was implemented. During heating and pressure increasing step, backbone carbon atoms (C*α*) were restrained using a spring force. The system was equilibrated for 50 ns at 300K and 1 bar without any restraint before moving to production run. A 2 fs timestep was used for the simulation. To stabilize the hydrogen bond vibration in the 2fs timestep, the SHAKE algorithm was used(*44*). A 10 *Å* cutoff was used for the nonbonded interactions. Long range electrostatic interactions are taken into account by the Particle Mesh Ewald method (*45*). Production runs of the simulation are performed in a NPT ensemble. Periodic boundary conditions are maintained throughout the simulation.

### Adaptive Sampling

Previous computational studies have shown that Na^+^binding to a GPCR is a slow process. With a single long trajectory, we may not able to observe the Na^+^binding. Therefore, to capture the Na^+^binding event, we implement adaptive sampling (*46*–*50*). Adaptive Sampling approaches have been successfully used to sample protein-ion binding(*33*), protein-ligand binding(*51, 52*), protein conformational change(*53*–*56*), folding(*57*) and protein-protein association(*58*). Adaptive sampling is an iterative sampling process to make simulation parallelizable by simultaneously running multiple short simulations. First, we project our simulation data onto reaction co-ordinates in which we want to make our sampling efficient. We then cluster the projected data into states using k-means clustering. We select the starting point for the next round of simulation from the least counted states. For this case, the reaction coordinates that are used to sample from our data include the distant between closest Na^+^ from D^2.50^. For CB_1_ and CB_2_, we perform *∼*37*µ*s and *∼*26*µ*s of aggregate simulation.

### Markov State Models

To capture the thermodynamics and kinetics information from simulation, we build Markov state model (MSM) on MD data. MSM assumes Markovian properties of MD data and accordingly generates ensemble distribution of the protein dynamic landscape (*59*–*61*). To build MSM describing the Na^+^ binding event, simulation data are represented using features (e.g. residue-residue distance, binding distance, dihedral angles) important to capture the binding process and important structural changes (Tables S1 and S2). For better approximation of MSM timescales, features was linearly transformed to time-independent components (tiCs) (*62, 63*). Tic helps to project the data along the slowest components for better characterization of slowest process (Figure S6). The projected space is then further discretized into states. MSM calculates the probability of the states and timescales of transition by estimating the eigenvectors and eigenvalues of the corresponding transition probability matrix, *T*. Each element of *T* (*T*_*ij*_), is estimated from the probability of jump between state i to state j at a particular lag time (*τ*). To find out the lag time at which the Markovian property is valid, implied timescale was calculated using Tics capturing 95 percent kinetic variance compared to all tic components and 30 ns of tic lag time. Minimum lag time at which implied time scale of slowest process converged (with 5 percent of two consecutive points) is selected to be the MSM lag time (Figures S20A and S20B). To optimize other hyperparameters (cluster numbers and number of tic components), VAMP2 scores are compared by building MSM with different cluster numbers and tic variational cutoff (Figures S20C and S20D) (*64*).

### Trajectory Analysis

MD trajectory features are calculated and analyzed using MDtraj (*65*) and CPPTRAJ (*66*). Pore tunnel radius calculation was performed using HOLE software (*67*). Pocket volume calculation was performed using Fpocket (*36*) and POVME (*68*). For trajectory visualization, VMD package is used (*69*).

### Standard Binding Free Energy Calculation

To calculate free energy, we project our data into x, y, and z component distance of Na^+^ from the D^2.50^ (*70*). The 3-D projection is clustered into into 300*300*300 bins. The standard binding free energy is calculated using the formula 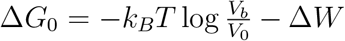 (*70*). In this case is the Δ W is the depth of 3-D potential mean force force calculated using MSM. *V*_0_ is equal to 1661 *Å*^3^ which is corresponds to 1M concentration of Na^+^. *V*_*b*_ is the weighted binding volume calculated using the formula *∫ exp*(*βW* (*x, y, z*))*dxdydz*.

### Transition Path Theory

Transition path theory (TPT) calculates the timescale for transition between different MSM macrostates by estimating mean free passage time. MFPT between macrostate A and B is determined by the equation where 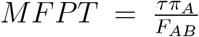 (*71*). where *π*_*A*_ is the probability of macrostate A calculated by MSM. *F*_*AB*_ is flux between macrostate A and B which is determined by the equation 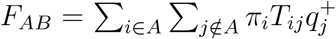 where 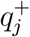 is probability of state j to move to B before A and *T*_*ij*_ is the probability of jump from state i to state j at lagtime *τ*. 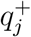 is estimated from the balance equation 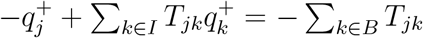. TPT calculation is performed using pyEMMA software package (*72*).

### Error Calculation

Errors on thermodynamics and kinetic calculations is determined by bootstrapping (*73*). We generate 200 rounds of bootstrap samples where each sample contains randomly picked 80% of total trajectories. We keep the state index same and build MSM for each of sample. Using the calculated stationary density and transition probability matrix of each sample, we determine error in our calculations.

