## Supplementary Information for "Distinct Binding Mechanisms for Allosteric Sodium Ion In Cannabinoid Receptors"

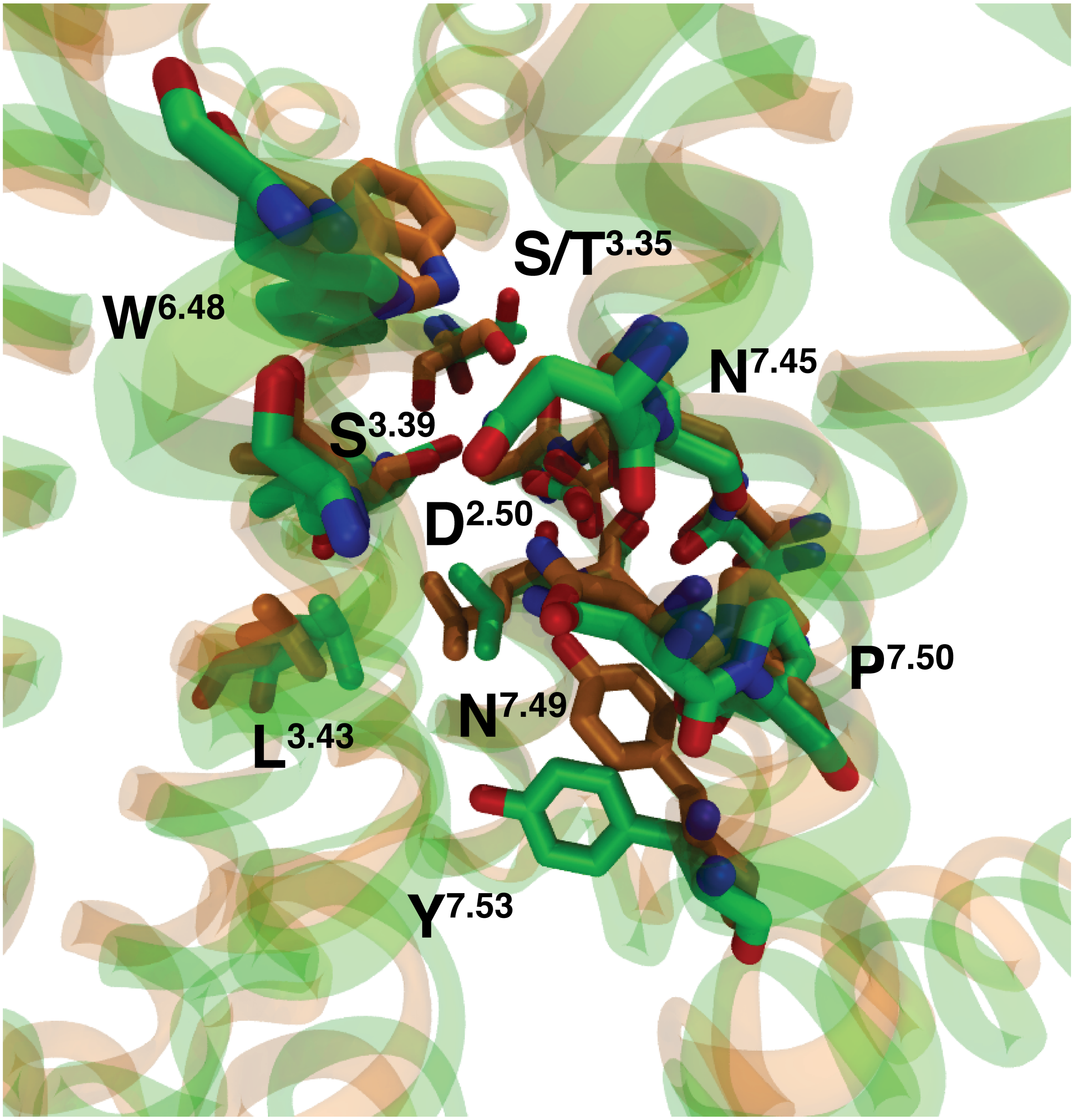


Figure S1: Superposition of inactive (PDB ID: 5TGZ, color: orange) $CB_1$ and inactive (PDB ID: 5TY, color: green) $CB_2$. $Na^+$ binding pocket residues are shown as sticks. Proteins are represented as cartoon.

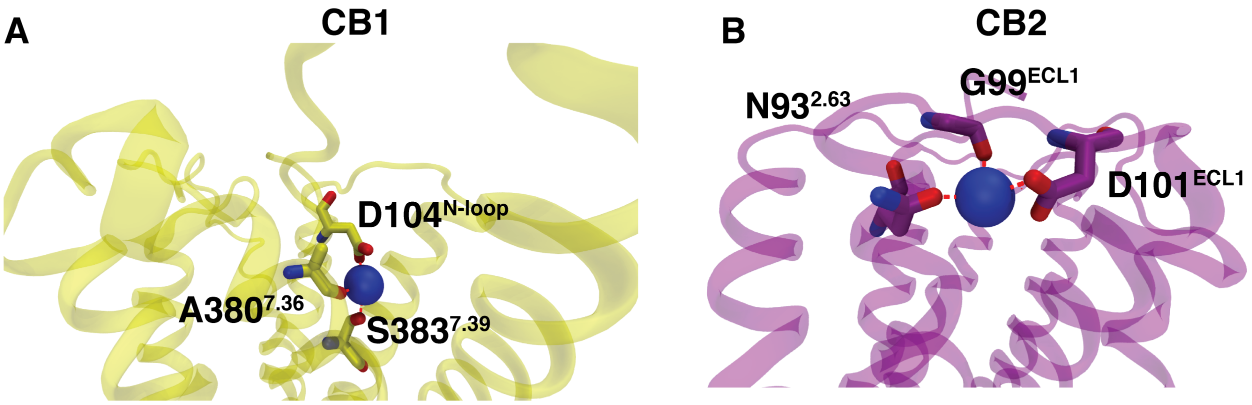


Figure S2: $Na^+$ interacting with residues in extracellular site. (A) shows a representative MD snapshot where $Na^+$ is interacting with polar residues in N-loop and orthosteric binding for $CB_1$. (B) shows a representative snapshot where $Na^+$ is ECL1 and TM1 residues for $CB_2$. Proteins are shown as Cartoon ($CB_1$: yellow, $CB_2$: voilet). Important residues are shown as sticks. $Na^+$s are represented as VDW representation (color: blue).

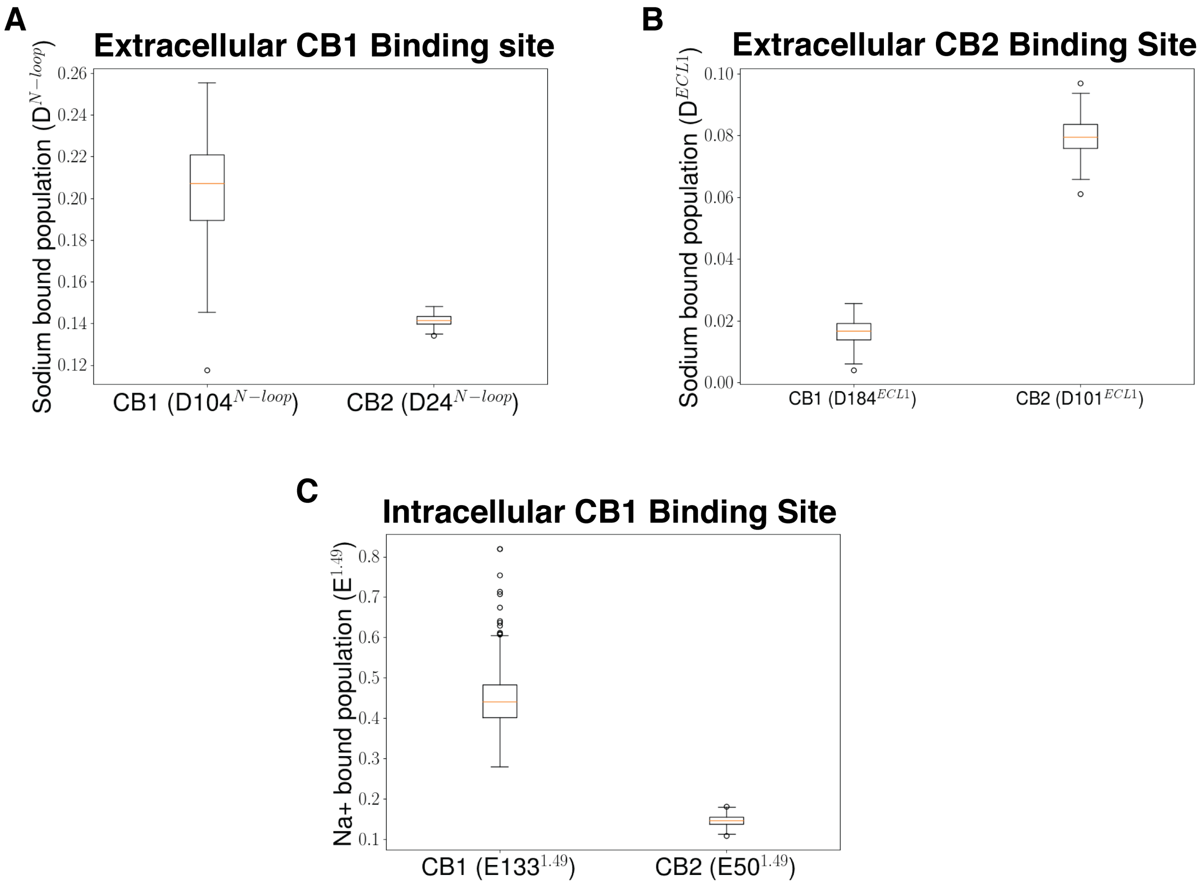


Figure S3: (A) Box plot shows distribution of fraction of $Na^+$ bound population for $CB_1$ and $CB_2$ in extracellular binding site for $CB_1$. (B) Box plot shows distribution of fraction of $Na^+$ bound population for $CB_1$ and $CB_2$ in extracellular binding site for $CB_2$. (C) Box plot shows distribution of fraction of $Na^+$ bound population for $CB_1$ and $CB_2$ in intracellular binding site for $CB_1$.Error calculations are performed with 200 bootstrap samples with 80% of total trajectories.

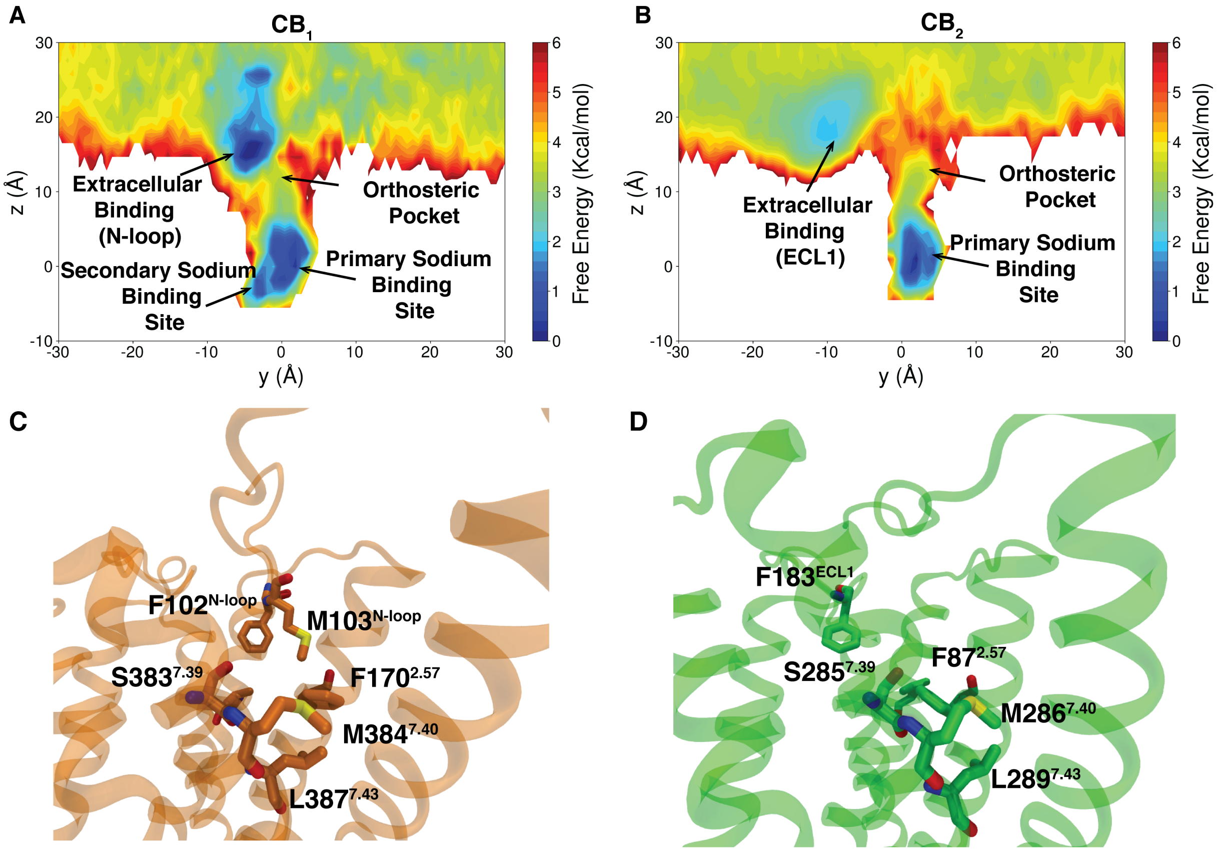


Figure S4: Extracellular $Na^+$ binding through orthosteric binding site. MSM weighted free energy landscapes projected along y and z component distance of $Na^+$ from D163$^{2.50}$ ($CB_1$) (A) and D80$^{2.50}$ ($CB_2$) (B). (C), (D) represent the residues in the orthosteric pocket along the $Na^+$ binding pathway for inactive $CB_1$ (PDB ID: 5TGZ) and $CB_2$ (PDB ID: 5ZTY). Proteins are shown as Cartoon ($CB_1$: orange, $CB_2$: green). Important residues are represented as sticks.

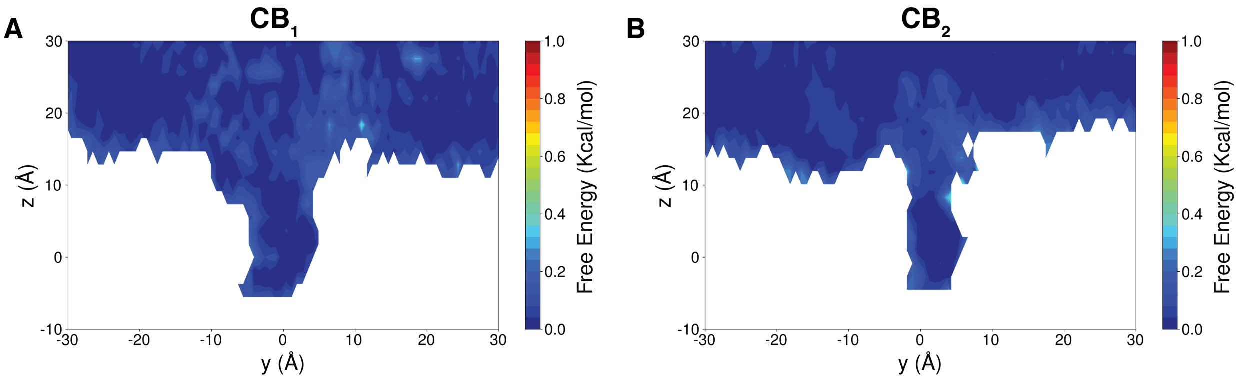


Figure S5: Error calculations on MSM weighted free energy landscapes projected along y and z component distance of $Na^+$ from D163$^{2.50}$ in CB$_1$ (A) and D80$^{2.50}$ in CB$_2$ (B). Error calculations are performed with 200 bootstrap samples with 80% of total trajectories.

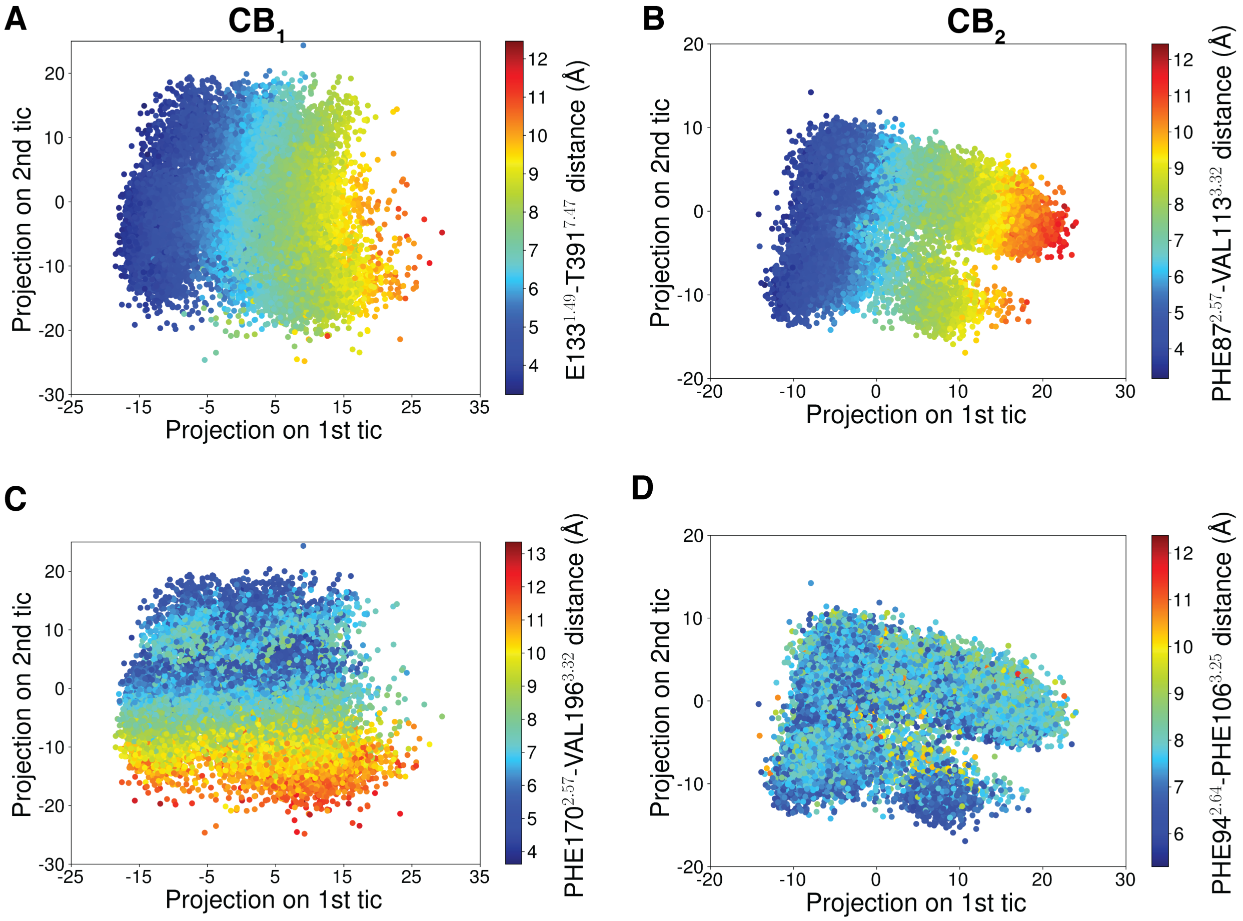


Figure S6: Variation of system features with highest correlation with tic 1 projected along tic 1 and tic 2 component for $CB_1$ (A) and $CB_2$ (B). Variation of system features with highest correlation with tic 2 are projected along tic 1 and tic 2 component for $CB_1$ (C) and $CB_2$ (D).

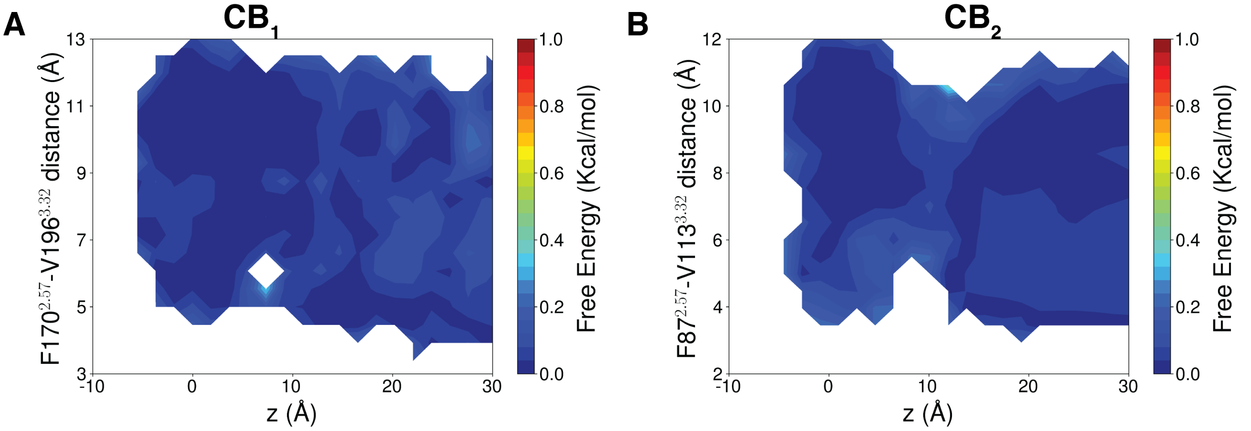


Figure S7: (A) Error calculation on MSM weighted free energy landscape projected along z component distance of $Na^+$ from D163$^{2.50}$ and F170$^{2.57}$ (C$\gamma$) -V196$^{3.32}$ (C$\gamma$) distance for $CB_1$. (B) Error calculations MSM weighted free energy landscape projected along z component distance of $Na^+$ from D80$^{2.50}$ and F87$^{2.57}$ (C$\gamma$) - V113$^{3.32}$ (C$\gamma$) distance for $CB_2$. Error calculations are performed with 200 bootstrap samples with 80% of total trajectories.

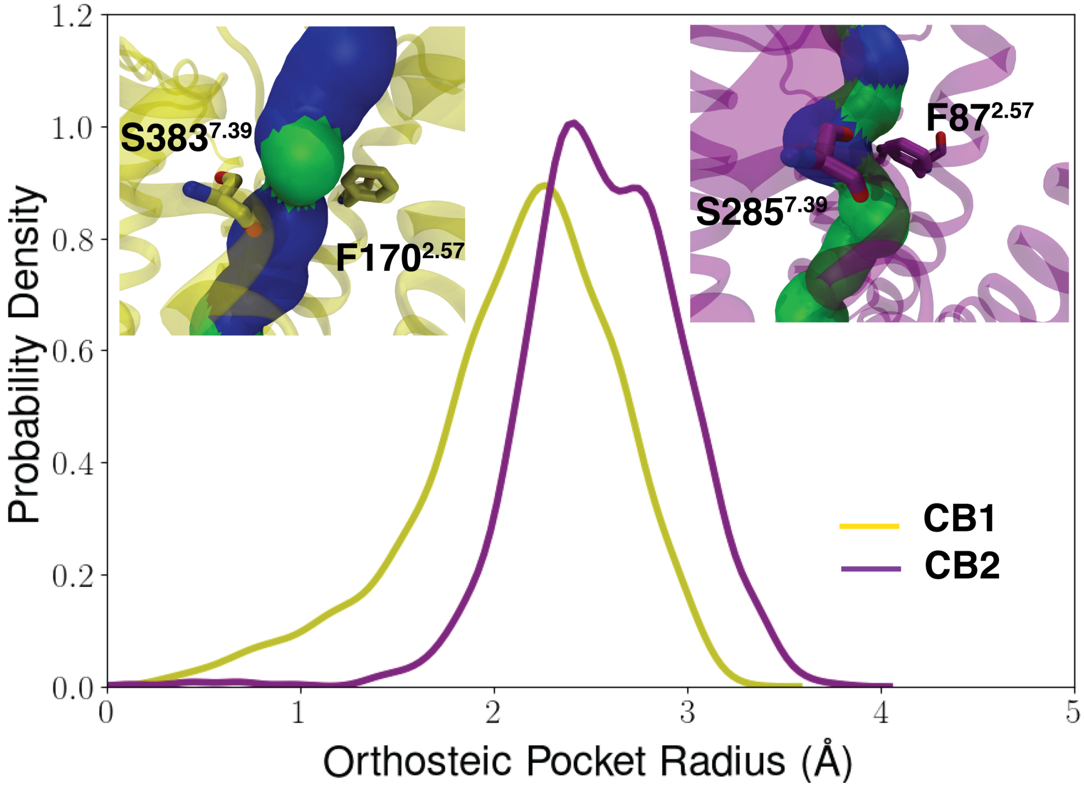


Figure S8: Distribution of radius in $Na^+$ binding tunnel in orthosteric pocket. Distributions are calculated with 1000 structures selected based on the MSM probability from the MSM microstates where $Na^+$ is bound in orthosteric binding pocket. Radius of the tunnel is calculated from the points where z cordinate lies between 0.5 Åin either side of average z cordinate between F170$^{2.57}$ (F87$^{2.57}$) , V196$^{3.32}$ (V113$^{3.32}$) and S383$^{7.39}$ (S285$^{7.39}$) for $CB_1$ ($CB_2$). Tunnel measurement is done using HOLE program. In inlet figures, proteins are shown as Cartoon ($CB_1$: yellow, $CB_2$: voilet). Important residues are represented as sticks.

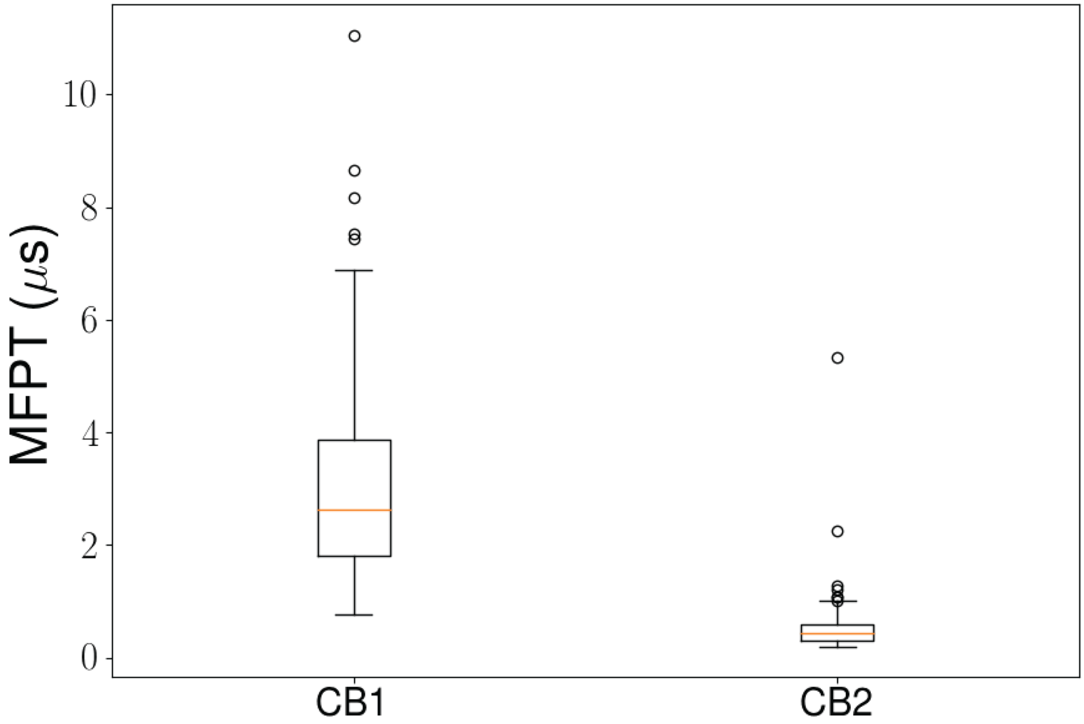


Figure S9: Box plot to show the distribution of timescale to jump from orthosteric pocket to primary $Na^+$ binding site. Timescale distributions are calculated by TPT analysis on 200 bootstrap samples with 80% of total trajectories.

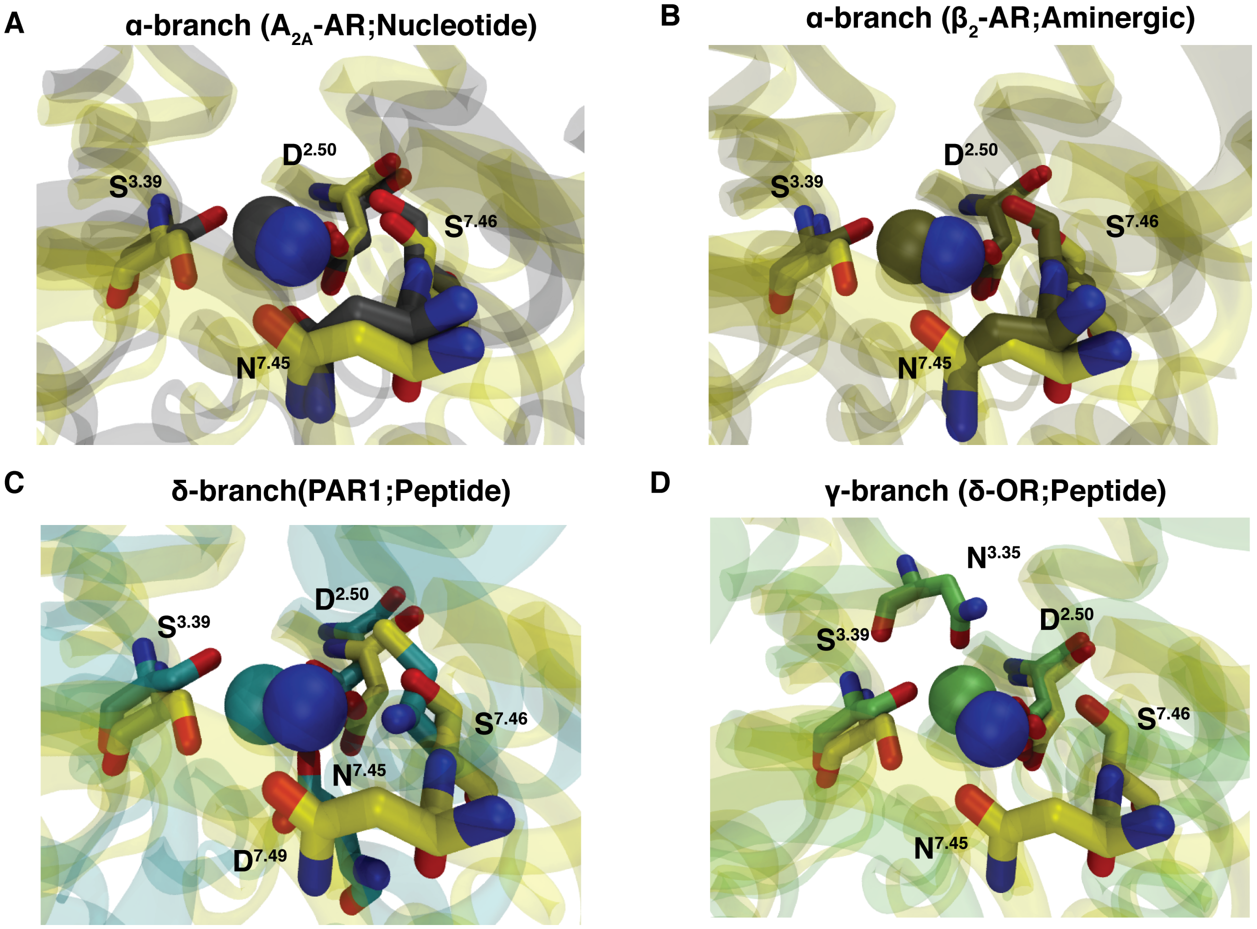


Figure S10: Comparison of $Na^+$ binding site of $CB_1$ with different branches of Class A GPCRs. (A) Residues within 4 Å of $Na^+$ are represented as sticks. Proteins are shown as Cartoon ($CB_1$: yellow, $A_{2A}$-AR: silver, $\beta_{2A}$-AR: tan, PAR1: cyan, $\delta$-OR: green).

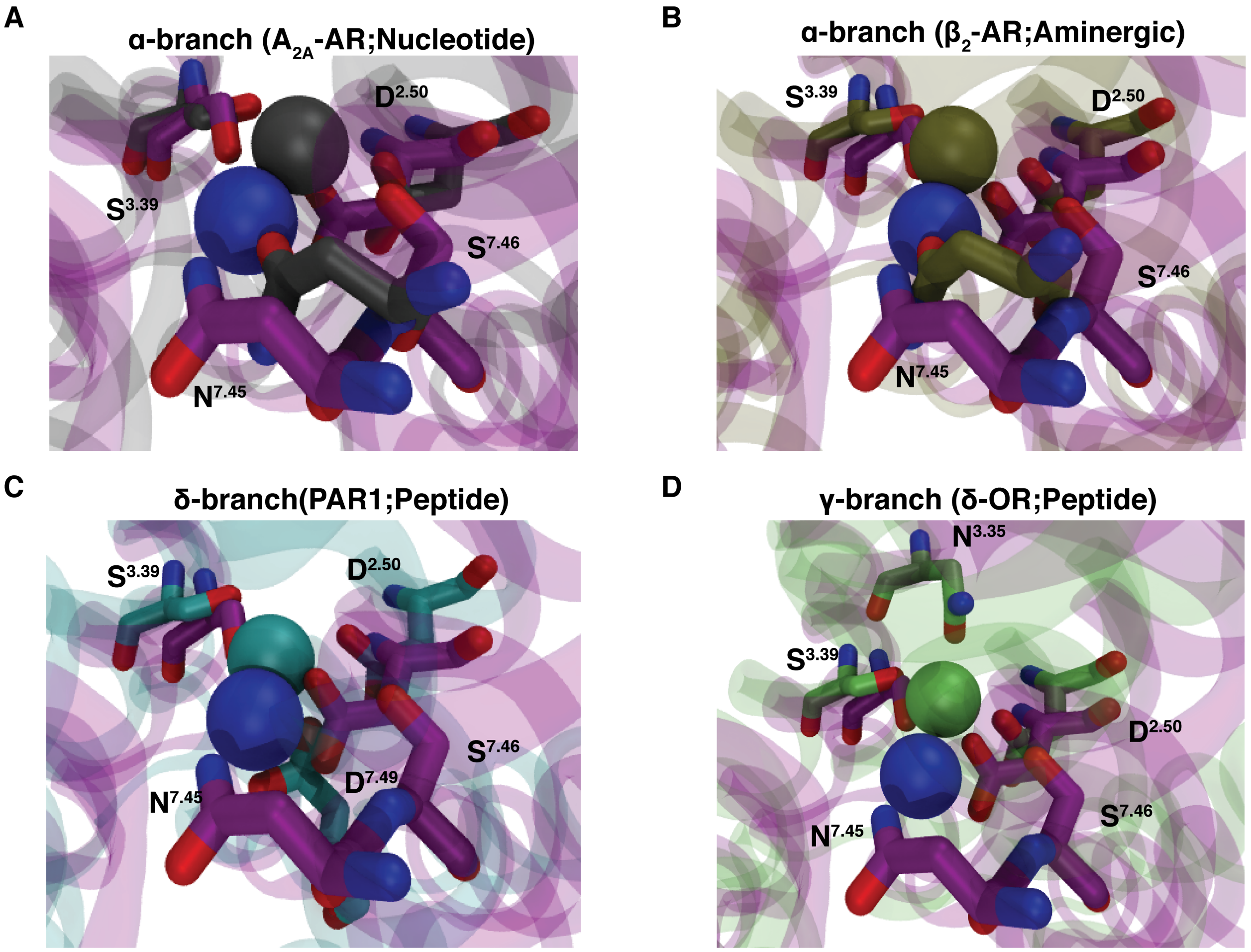


Figure S11: Comparison of $Na^+$ binding site of $CB_2$ with different branches of Class A GPCRs. (A) Residues within 4 $\mathring{A}$ of $Na^+$ are represented as sticks. Proteins are shown as Cartoon ($CB_2$: violet, $A_{2A}$-AR: silver, $\beta_{2A}$-AR: tan, PAR1: cyan, $\delta$-OR: green).

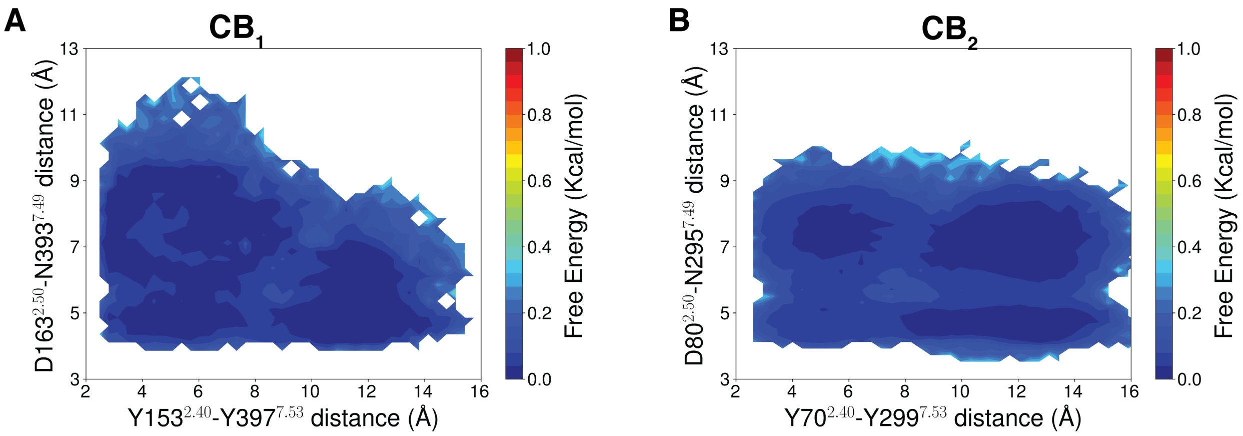


Figure S12: (A) Error calculations on MSM weighted free energy landscape projected along $Y153^{2.40}$-$Y397^{7.53}$ distance and $D163^{2.50}$-$N393^{7.49}$ distance for $CB_1$. (B) Error calculations on MSM weighted free energy landscape projected along $Y70^{2.40}$-$Y299^{7.53}$ distance and $D80^{2.50}$-$N295^{7.49}$ distance for $CB_2$. Error calculations are performed with 200 bootstrap samples with 80% of total trajectories.

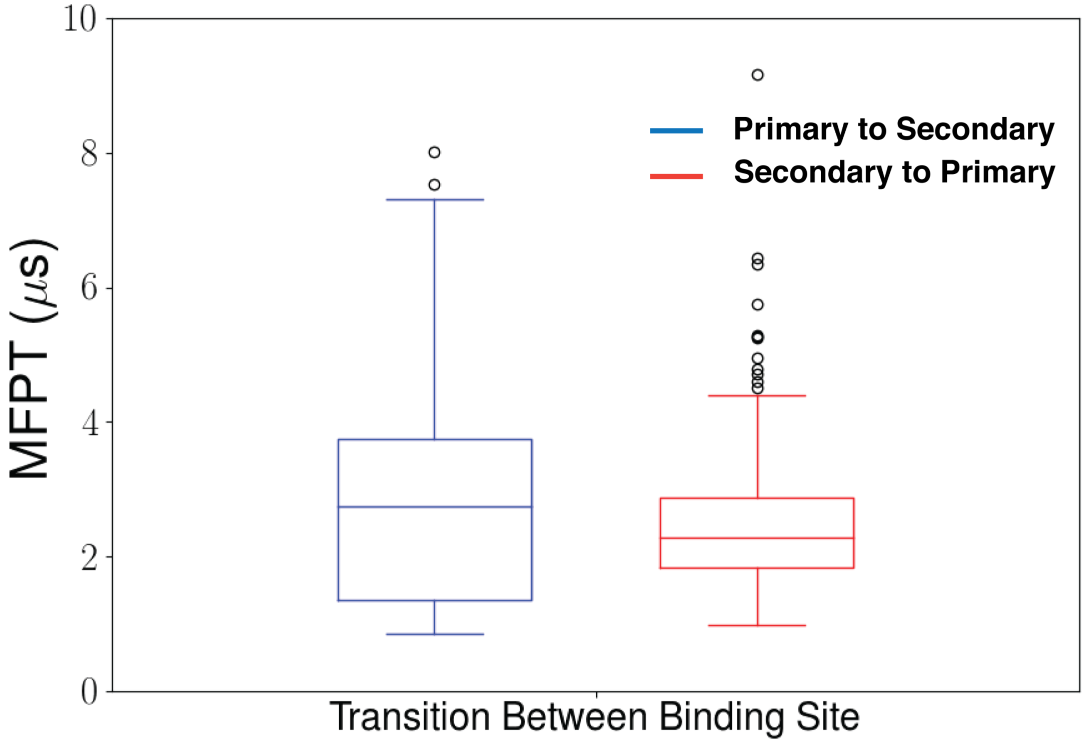


Figure S13: Box plot to show the distribution of timescale to jump from primary $Na^+$ binding site to secondary binding site and vicevarsa for $CB_1$. Timescale distributions are calculated by TPT analysis on 200 bootstrap samples with 80% of total trajectories.

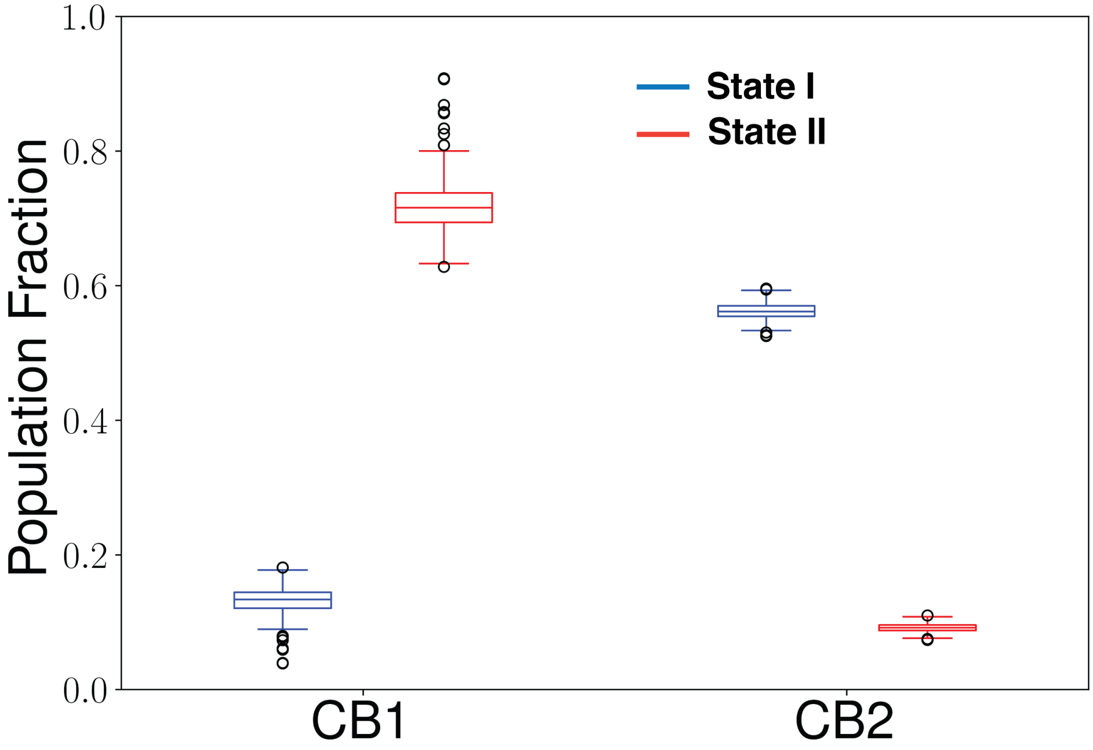


Figure S14: Box plot to show the distribution of population in state I (color: Blue) and state II (color: Red) for $CB_1$ and $CB_2$, respectively. Population is calculated using 200 bootstrap samples with 80% of total trajectories.

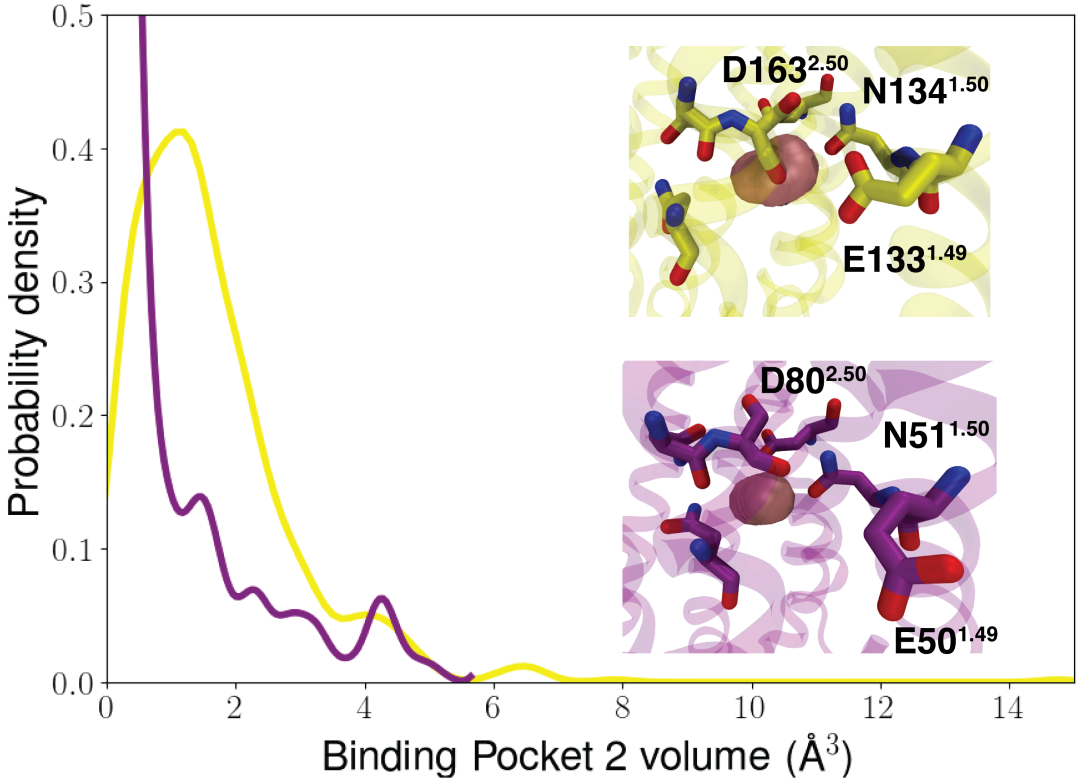


Figure S15: Volume distribution of secondary $Na^+$ binding site for $CB_1$ and $CB_2$. In inlet picture, MD snapshots with mean binding volume are represented. Calculated volumes are represented as surface. Residues surrounding secondary $Na^+$ binding site are shown as sticks.Proteins are shown as Cartoon ($CB_1$: yellow, $CB_2$: voilet).

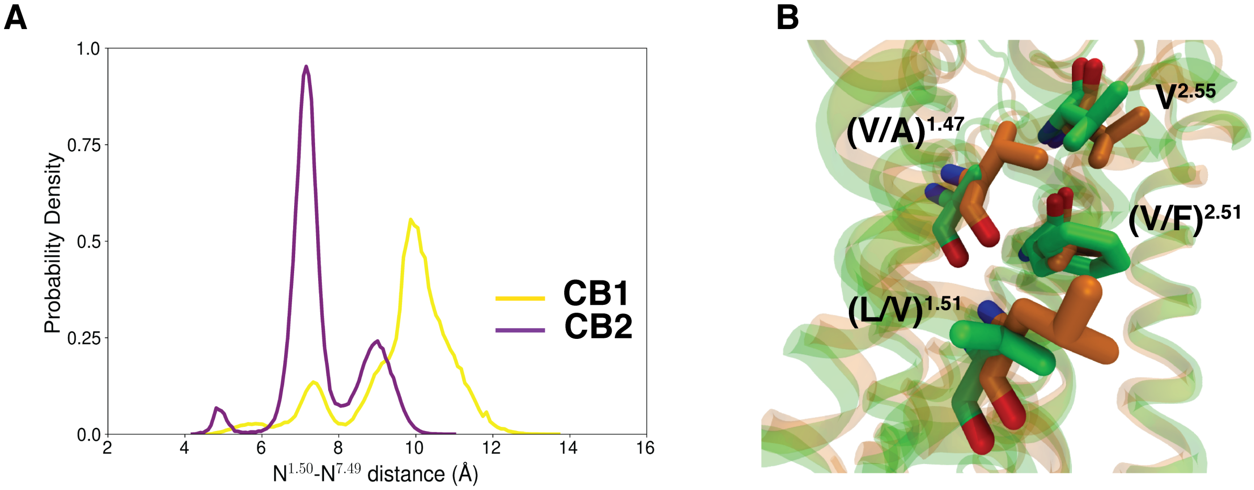


Figure S16: (A) MSM weighted distance distribution between $N^{1.50}$ and $N^{7.50}$. (B) Superposition of inactive structures of $CB_1$ and $CB_2$ from membrane side to focus on residues in TM1 and TM2 close to $N^{1.50}$. Proteins are shown as Cartoon ($CB_1$: yellow, $CB_2$: voilet). Important residues are represented as sticks.

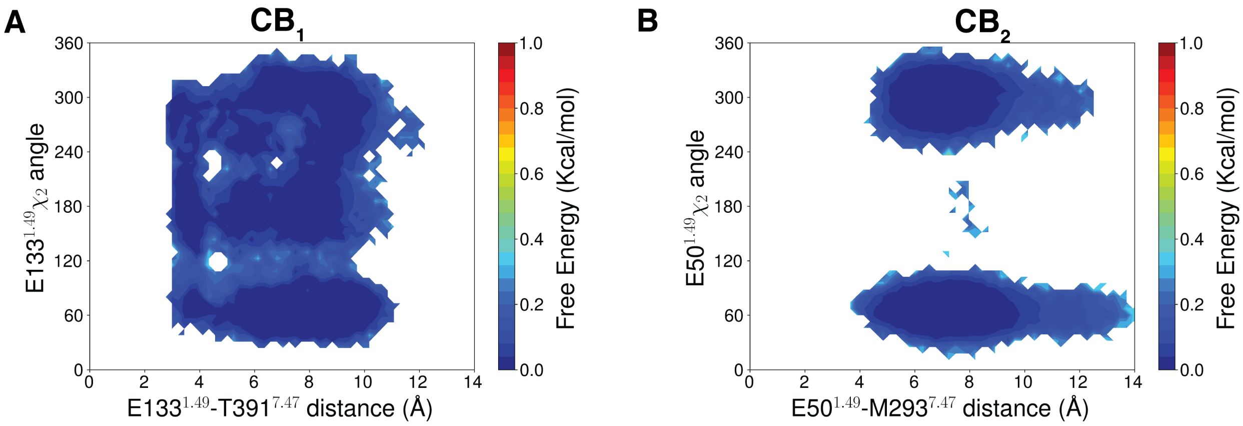


Figure S17: (A) Error calculations on MSM weighted free energy landscape between E133$^{1.49}$-T391$^{7.47}$ distance and E133$^{1.49}$ sidechain $\chi_2$ anagle for CB$_1$. (B) Error calculations on MSM weighted free energy landscape between E50$^{1.49}$-M293$^{7.47}$ distance and E50$^{1.49}$ sidechain $\chi_2$ angle for CB$_1$. Error calculations are performed with 200 bootstrap samples with 80% of total trajectories.

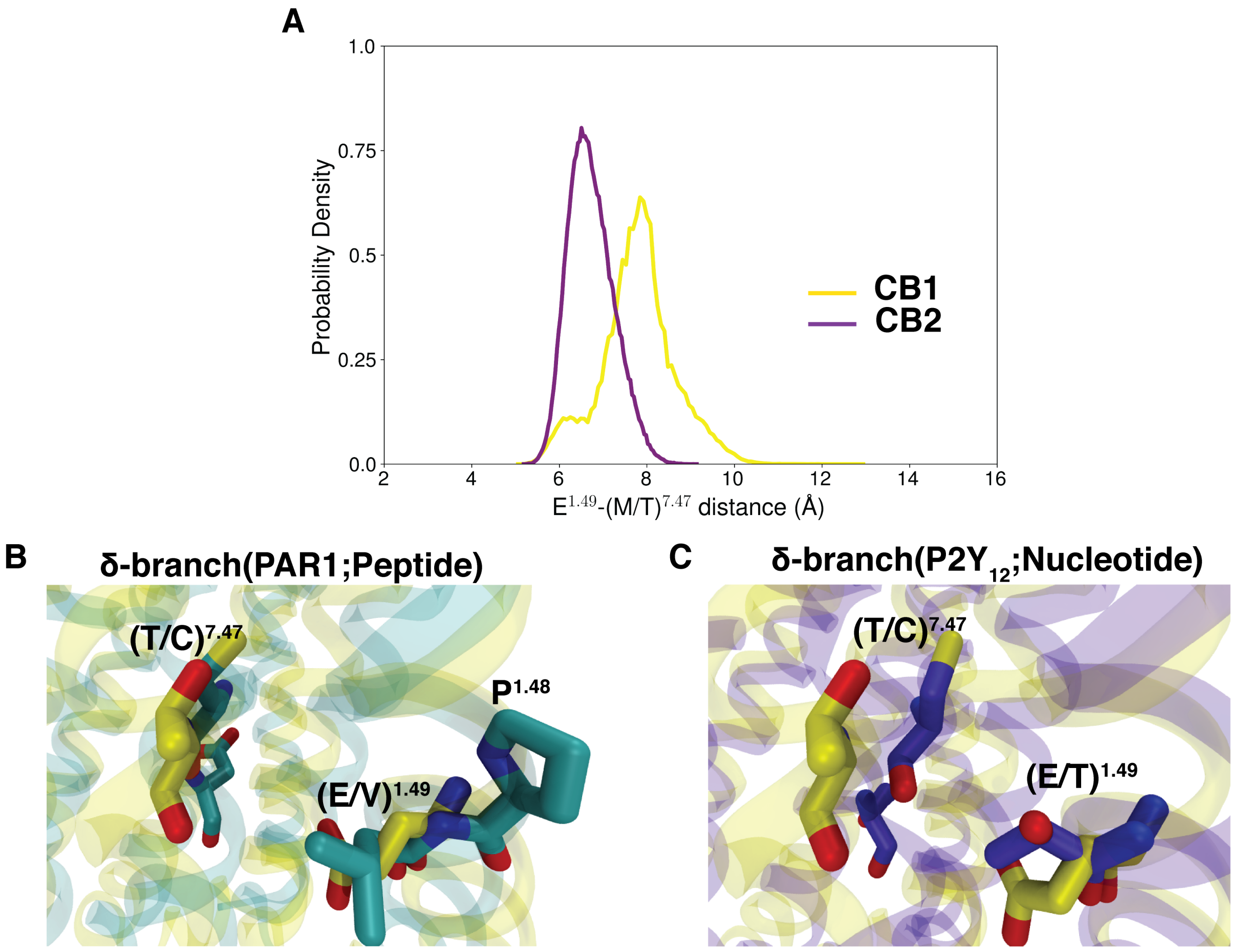


Figure S18: (A) MSM weighted distance distribution between E and T. (B) Superposition of inactive structures of $CB_1$ and PAR1 from membrane side to focus on residues in TM1 and TM7 close to intracellular binding side. (C) Superposition of inactive structures of $CB_1$ and P2Y12 from membrane side to focus on residues in TM1 and TM7 close to intracellular binding side. Proteins are shown as Cartoon ($CB_1$: yellow, PAR1: cyan, P2Y12: blue). Important residues are represented as sticks.

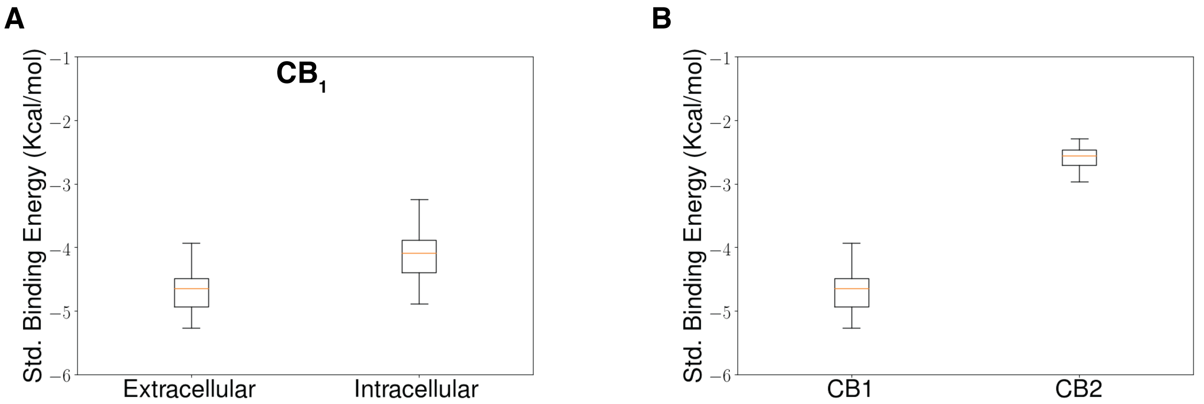


Figure S19: (A) Box plot to show the distribution of standard binding free energy for extracellular and intracellular binding of $Na^+$ for $CB_1$. (B) Box plot to show the distribution of standard binding free energy for extracellular binding for $CB_1$ and $CB_2$. Energy distributions are calculated from 20 bootstrap samples with 80% of total trajectories.

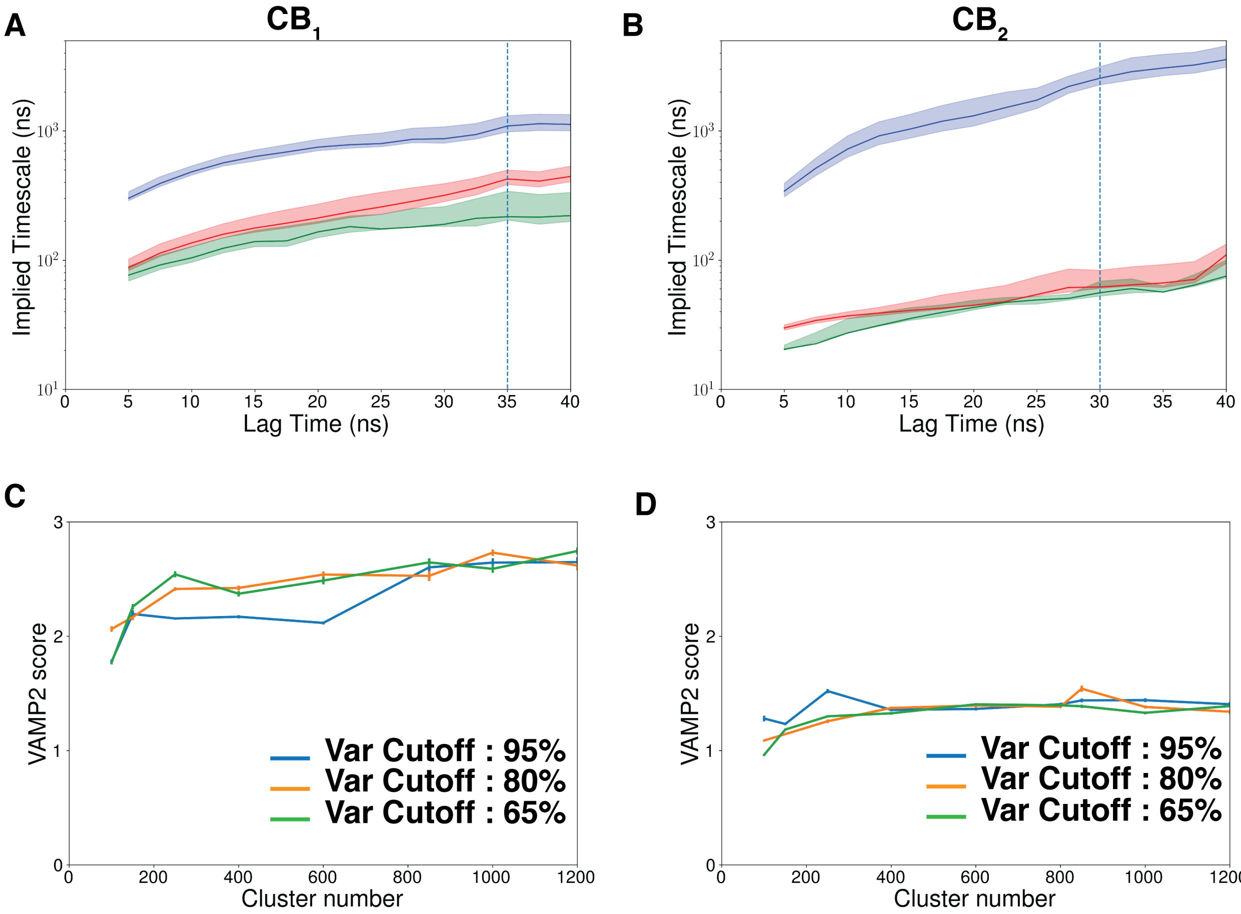


Figure S20: The timescales calculated from the 2nd, 3rd and 4th highest MSM eigenvalues for $CB_1$ (A) and $CB_2$ (B). VAMP-2 scores with different clusters and different tica variance for $CB_1$ (C) and $CB_2$ (D). MSM lagtime of 35ns and 30ns was chosen for $CB_1$ and $CB_2$, respectively.

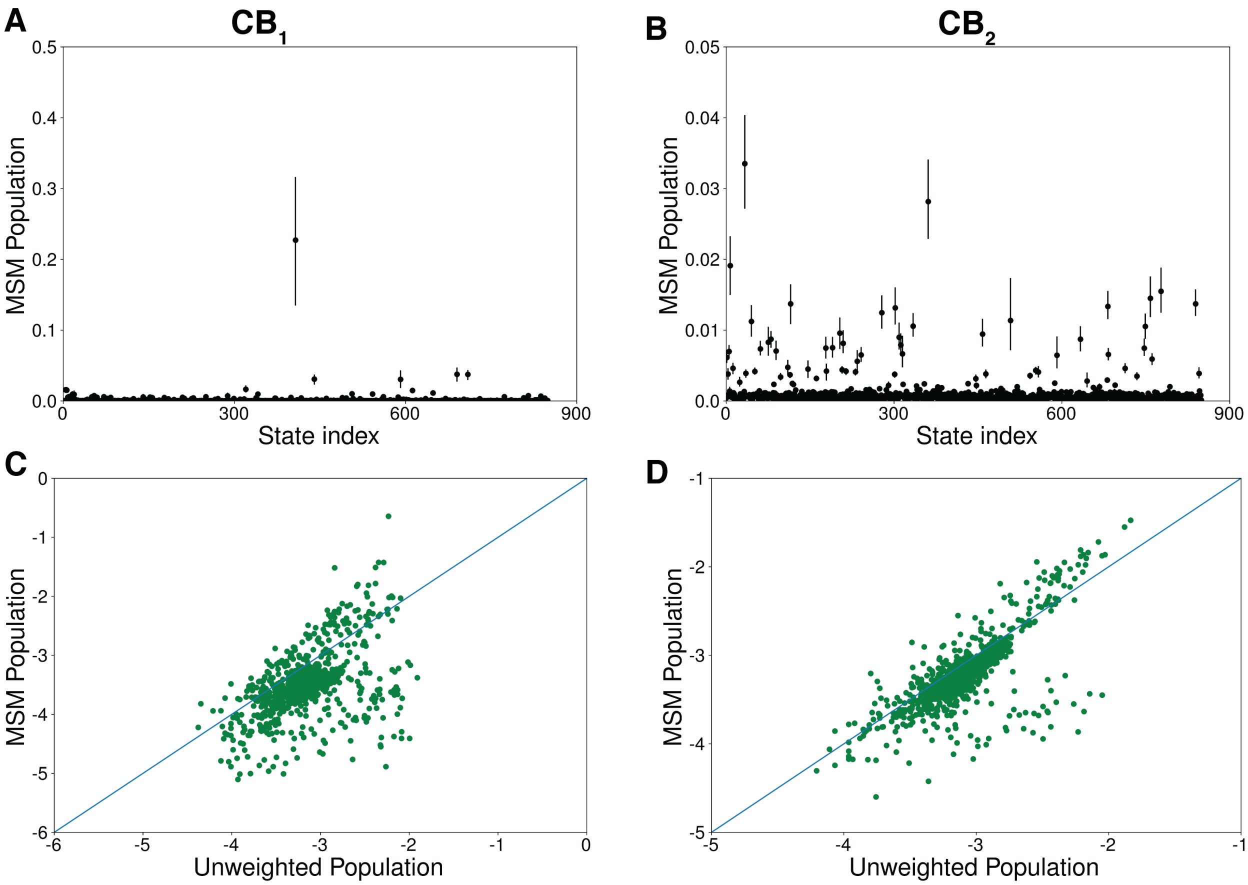


Figure S21: MSM population density for different MSM state index for $CB_1$ (A) and $CB_2$ (B). Total count variation in unweighted and MSM weighted states for $CB_1$ (C) and $CB_2$ (D). Error calculations are performed with 200 bootstrap samples with 80% of total trajectories.

Table S1: Distance and angle features used to build Markov state model for $CB_1$.

| Position | Number | Type | Feature |
|---|---|---|---|
| $Na^+$ binding | 1 | distance from $D163^{2.50}$ | x-component distance of closest $Na^+$ |
| | 2 | distance from $D163^{2.50}$ | y-component distance of closest $Na^+$ |
| | 3 | distance from $D163^{2.50}$ | z-component distance of closest $Na^+$ |
| Residue movement along the pathway | 4 | distance | $F170^{2.57}(C\gamma)$-$V196^{3.32}(C\gamma)$ |
| | 5 | distance | $T130^{1.46}(C\gamma)$-$L387^{7.43}(C\gamma)$ |
| | 6 | distance | $N134^{1.50}(C\gamma)$-$D163^{2.50}(C\gamma)$ |
| | 7 | distance | $E133^{1.49}(C\gamma)$-$T391^{7.47}(C\gamma)$ |
| | 8 | Dihedral Angle ($\chi_1$) | $E133^{1.49}$ |
| | 9 | Dihedral Angle ($\chi_1$) | $N134^{1.50}$ |
| | 10 | Dihedral Angle ($\chi_1$) | $D163^{2.50}$ |
| | 11 | Dihedral Angle ($\chi_1$) | $F170^{2.57}$ |
| | 12 | Dihedral Angle ($\chi_1$) | $L387^{7.43}$ |
| | 13 | Dihedral Angle ($\chi_1$) | $N389^{7.45}$ |

Table S2: Distance and angle features used to build Markov state model for $CB_2$.

| Position | Number | Type | Feature |
|---|---|---|---|
| $Na^+$ binding | 1 | distance from $D80^{2.50}$ | x-component distance of closest $Na^+$ |
| | 2 | distance from $D80^{2.50}$ | y-component distance of closest $Na^+$ |
| | 3 | distance from $D80^{2.50}$ | z-component distance of closest $Na^+$ |
| Residue movement along the pathway | 4 | distance | $F87^{2.57}(C\gamma)$-$V113^{3.32}(C\gamma)$ |
| | 5 | distance | $F91^{2.61}(C\gamma)$-$I110^{3.29}(C\gamma)$ |
| | 6 | distance | $F94^{2.64}(C\gamma)$-$F106^{3.25}(C\gamma)$ |
| | 7 | Dihedral Angle ($\chi_1$) | $D80^{2.50}$ |
| | 8 | Dihedral Angle ($\chi_1$) | $F87^{2.57}$ |
| | 9 | Dihedral Angle ($\chi_1$) | $F94^{2.64}$ |
| | 10 | Dihedral Angle ($\chi_1$) | $I110^{3.29}$ |
| | 11 | Dihedral Angle ($\chi_1$) | $N291^{7.45}$ |
